# Genetic dissection of the mitochondrial lipoylation pathway in yeast

**DOI:** 10.1101/2020.11.24.395780

**Authors:** Laura P. Pietikäinen, M. Tanvir Rahman, J. Kalervo Hiltunen, Carol L. Dieckmann, Alexander J. Kastaniotis

**Affiliations:** Faculty of Biochemistry and Molecular Medicine and Biocenter Oulu, University of Oulu, PO Box 5400, Oulu FI-90014, Finland; Department of Molecular and Cellular Biology, University of Arizona, Tucson, AZ 85721, USA

**Keywords:** lipoylation, mitochondrial fatty acid synthesis (mtFAS), octanoyl/lipoyl transferases, *S. cerevisiae* model, supplementation studies, Lip3/LIPT1, Lip2/LIPT2, Lip3 substrate, lipoylation disorders

## Abstract

**Background:** Lipoylation of 2-ketoacid dehydrogenases is essential for mitochondrial function in eukaryotes. While the basic principles of the lipoylation processes have been worked out, we still lack a thorough understanding of the details of this important post-translational modification pathway. Here we used yeast as a model organism to characterize substrate usage by the highly conserved eukaryotic octanoyl/lipoyl transferases *in vivo* and queried how amenable the lipoylation system is to supplementation with exogenous substrate.

**Results:** We show that the requirement for mitochondrial fatty acid synthesis to provide substrates for lipoylation of the 2-ketoacid dehydrogenases can be bypassed by supplying the cells with free lipoic acid (LA) or octanoic acid (C8) and a mitochondrially targeted fatty acyl/lipoyl activating enzyme. We also provide evidence that the *S. cerevisiae* lipoyl transferase Lip3, in addition to transferring LA from the glycine cleavage system H protein to the pyruvate dehydrogenase (PDH) and α-ketoglutarate dehydrogenase (KGD) E2 subunits, can transfer this cofactor from the PDH complex to the KGD complex. In support of yeast as a model system for human metabolism, we demonstrate that the human octanoyl/lipoyl transferases can substitute for their counterparts in yeast to support respiratory growth and protein lipoylation. Like the wild-type yeast enzyme, the human lipoyl transferase LIPT1 responds to LA supplementation in the presence of the activating enzyme LplA.

**Conclusions:** In the yeast model system, the eukaryotic lipoylation pathway can use free LA and C8 as substrates when fatty/lipoic acid activating enzymes are targeted to mitochondria. Lip3 LA transferase has a wider substrate specificity than previously recognized. We show that these features of the lipoylation mechanism in yeast are conserved in mammalian mitochondria. Our findings have important implications for the development of effective therapies for the treatment of LA or mtFAS deficiency-related disorders.

## BACKGROUND

Like its functional cousin biotin, the enzyme cofactor lipoic acid (LA) acts as a “swinging arm” moiety to shuttle reaction intermediates from one enzymatic reaction center to another in multidomain protein complexes catalyzing oxidative decarboxylation of 2-ketoacids [1]. In eukaryotes, the octanoic acid (C8) precursor of endogenously synthesized LA is generated by mitochondrial fatty acid synthesis (mtFAS). In humans, all LA-dependent enzyme complexes are mitochondrial: pyruvate dehydrogenase, (PDH), α-ketoglutarate dehydrogenase (KGD), the glycine cleavage system (GCS), branched chain dehydrogenase (BCD), and α-ketoadipate dehydrogenase (OAD) [2].

With the exceptions of BCD and OAD, the LA-dependent complexes are found also in mitochondria of the yeast *Saccharomyces cerevisiae*. Invariantly, LA is attached to the N^6^-amino group of specific lysine residues of the E2 (PDH, KGD, BCD) or H (GCS) protein subunits of these complexes in a stable amide linkage. Because of the central role for KGD in the tricarboxylic acid cycle, LA is essential for cells and organisms that cannot satisfy their energy demands solely by glycolysis. Although decades have passed since the discovery and characterization of LA, and the essentials of synthesis and attachment of LA in *Escherichia coli* and *Bacillus subtilis* have been worked out, our understanding of the basic mechanism of lipoylation in eukaryotes (Figure 1) and the substrates used in these processes has improved only recently [3, 4].

**Figure 1.**
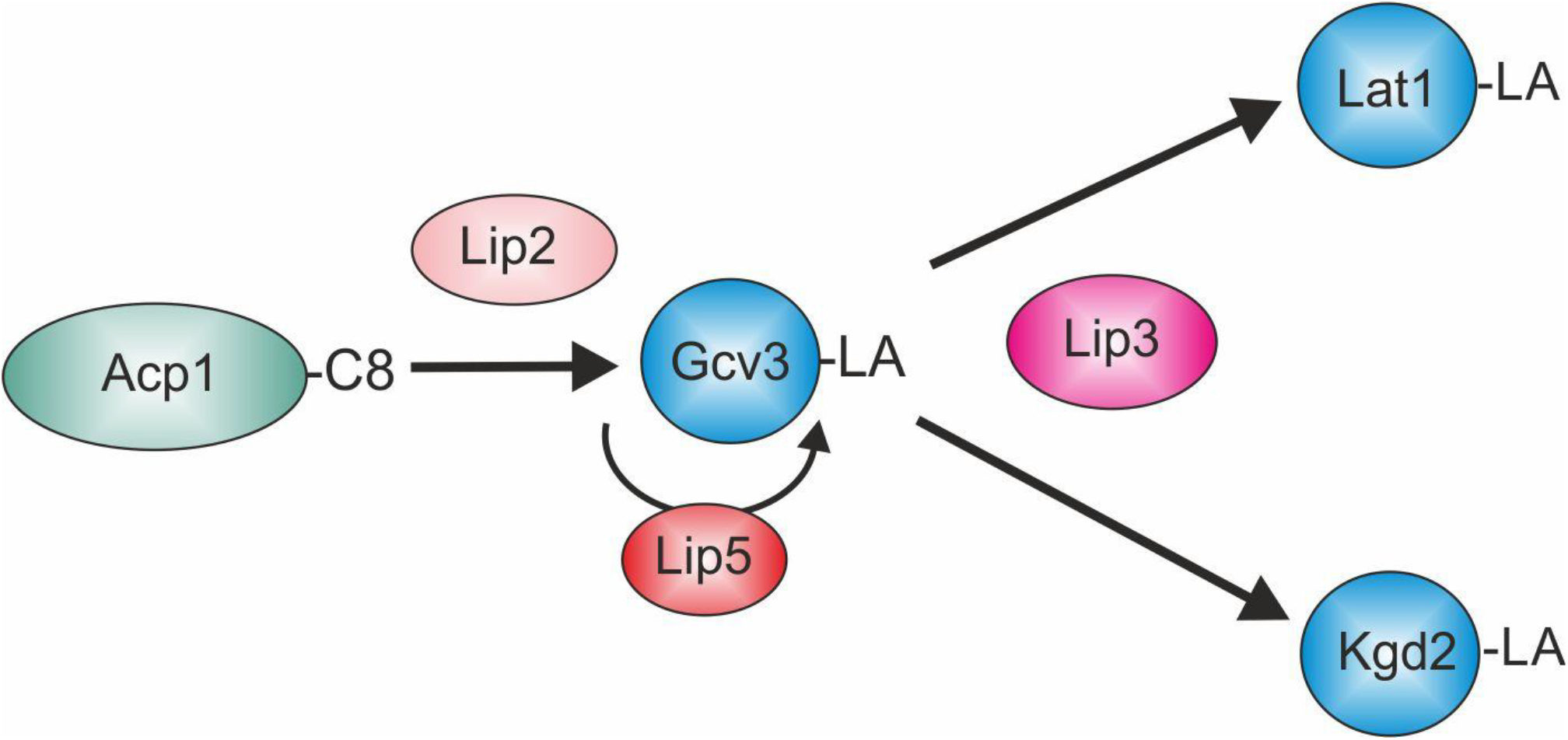
Cartoon depicting the current model of the lipoylation process in eukaryotic mitochondria. Octanoylated ACP generated by mtFAS serves as a substrate for transfer of the C8 moiety by the octanoyl transferase Lip2 (LIPT2 in humans) to Gcv3, the H protein of the GCS (GCSH). The octanoyl moiety is then converted to the lipoyl cofactor by lipoyl synthase Lip5 (LIAS), and subsequently transferred to the E2 subunits of other ketoacid dehydrogenase complexes (depicted are Lat1 and Kgd2 of the *S. cerevisiae* PDH and KGD complexes) by lipoyl transferase Lip3 (LIPT1).

A basic set of three enzymes appears to be required for lipoylation: lipoyl synthase LipA/Lip5/LIAS (*E. coli/S. cerevisiae/*humans) [5–7], and the lipoyl/octanoyl transferases LipB/Lip2/LIPT2 [7–9] and LplA/Lip3/LIPT1 [10–13]. In *E. coli*, the LipB transferase in combination with the LA synthase LipA, is sufficient for lipoylation of all LA-requiring enzymes. In an alternative route for protein lipoylation in *E. coli*, the lipoyl transferase LplA activates exogenously available free LA by adenylation, and then transfers it to the target proteins. In yeast, Lip2 can modify only the Gcv3 H-subunit of the GCS. Lipoylated Gcv3 is then the substrate for subsequent lipoylation of the E2 subunits by Lip3-catalyzed transamidation [11]. There is no evidence for the existence of a native *S. cerevisiae* LA-scavenging pathway analogous to the *E. coli* LplA– dependent route, since LA supplementation does not improve growth of lipoylation- or mtFAS-deficient yeast strains on non-fermentable carbon sources, and mtFAS appears to be the sole producer of the C8 used as a precursor for LA synthesis in mitochondria [6, 9, 14]. The schematic depicted in Figure 1 summarizes the current model of the lipoylation pathway in eukaryotes.

Several groups of disorders caused by of inborn errors leading in humans that affect lipoylation in mitochondria have been described [15]. The first group encompasses defects due to failures in synthesis of Fe-S clusters [16–19]. A second group of mitochondrial diseases is defined by mutations in lipoic acid synthase, Lip5/LIAS, (*S. cerevisiae*/humans) and protein lipolylation defects [7, 12, 16, 20]. A third group includes mtFAS disorders such as MCAT (malonyl-CoA transferase) deficiency [21] and MEPAN (mitochondrial enoyl-CoA reductase protein associated neurodegeneration) [22], the latter of which leads to mitochondrial enzyme complex hypolipoylation. In mice, deletion of the *Mecr* gene encoding enoyl reductase, the last enzyme in the mtFAS pathway, leads to loss of lipoylated proteins and early embryonic death [23].

Unlike yeast, mammalian mitochondria harbor fatty acyl activating enzymes, which raises the possibility that mtFAS or lipoylation pathway disorders may be alleviated by nutritional supplementation. To address this issue in a simple eukaryotic model, we set out to investigate whether ectopic expression of LA- or fatty acid-activating enzymes in yeast mitochondria combined with addition of LA or C8 to the growth media would rescue lipoylation and/or growth defects of mutant strains. Expanding on this goal, we generated tools that allowed us to dissect individual steps of the lipoylation pathway in more detail, including mitochondrially localized variants of LplA and Fam1-1 acyl-CoA ligases. We observed that exogenous LA/fatty acids are available for lipoylation processes in yeast mitochondria when activating enzymes are engineered to localize to the organelle. We also provide evidence that Lip3 acts as a multidirectional lipoyl transamidase in yeast mitochondria. Finally, we demonstrate that human lipoyl transferases can functionally replace their yeast homologs. Our work provides novel insights into the eukaryotic lipoylation pathway, suggesting potential routes to development of effective treatments for mtFAS-related disease.

## RESULTS

### Mitochondrially localized Fam1-1 acyl-CoA ligase plus C8 or LA supplementation bypasses mtFAS and rescues lipoylation

Yeast strains harboring mtFAS loss-of-function mutations are respiratory deficient due to defects in protein lipoylation, RNA processing, and assembly of the OXPHOS-complexes [11, 24, 25]. To investigate whether fatty acid substrates fed to yeast, can be transported into mitochondria and activated by a purposefully mislocalized enzyme, we took advantage of the *FAM1-1* suppressor allele which encodes a mitochondrially mislocalized fatty acyl-CoA ligase that restores the growth of mtFAS mutant strains on nonfermentable carbon sources [26]. In previous studies that used this mitochondrially mislocalized fatty acyl-CoA ligase [26–29], growth of cells deleted for the mtFAS 3-hydroxyacyl thioester dehydrogenase of mtFAS (Δ*htd2*), as well as strains deleted for all other mtFAS enzymes, was restored on unsupplemented synthetic media containing glycerol as carbon source. However, we and others have observed that lipoylation of ketoacid dehydrogenase E2 subunits was not detected upon *FAM1-1* suppression [27, 28].

We reasoned that one possible explanation for the lack of restoration of lipoylation in mtFAS mutants by Fam1-1 is that free C8 is not available in mitochondria for activation and as substrate for LA synthesis. Thus, we investigated whether protein lipoylation could be rescued in mtFAS-deficient yeast cells by growing strains harboring the *FAM1-1* allele on either minimal synthetic complete glycerol plates (SCG) or on rich glycerol plates (YPG) supplemented with LA or C8 (Figure 2).

**Figure 2.**
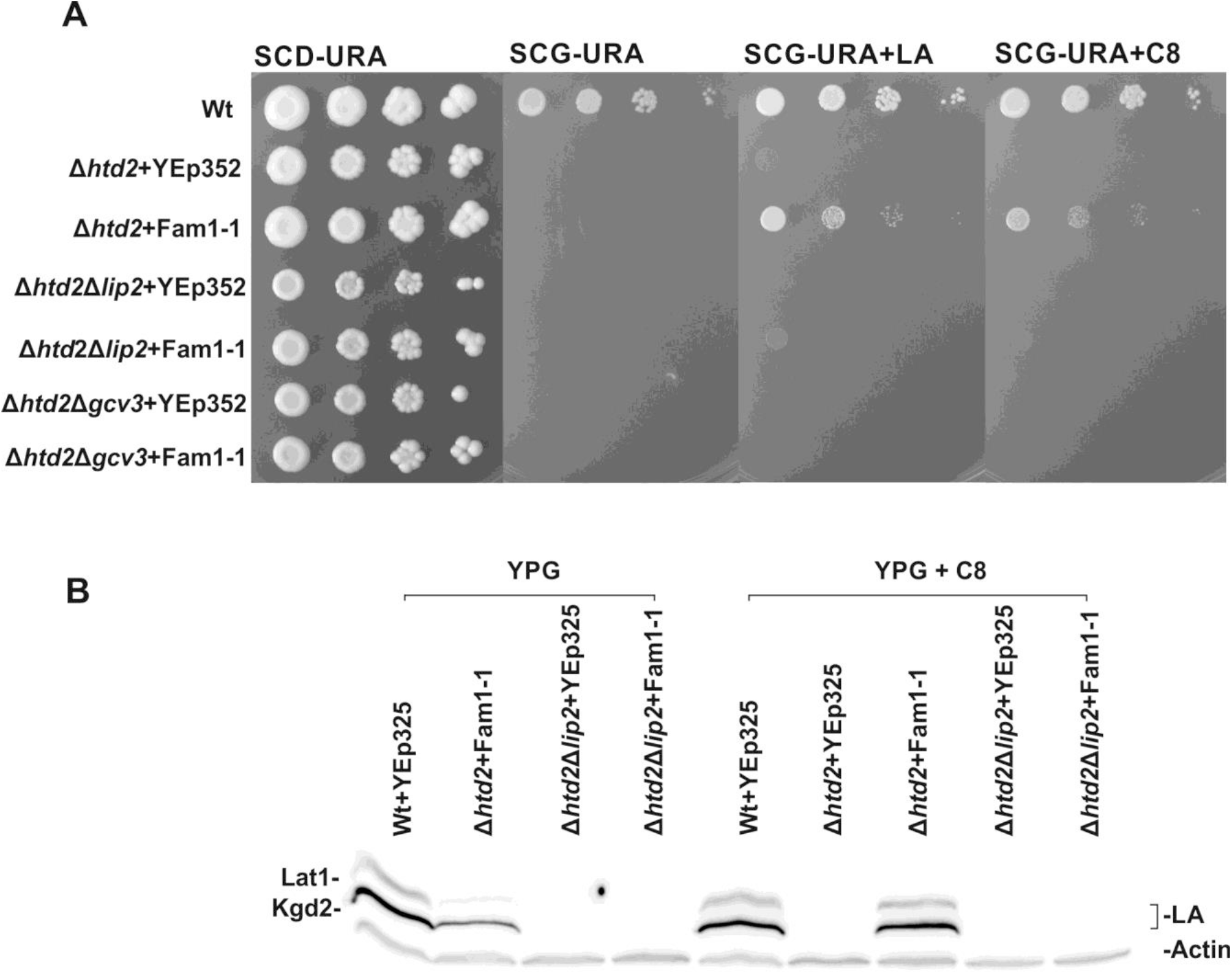
Analysis of lipoylation deficient strains expressing Fam1-1 (mitochondrially targeted Fam1/Faa2). All plasmids transformed into the different strains are based on the YEp352 yeast episomal plasmid backbone. (A) Growth assays of wild type +Yep352, Δ*htd2*+YEp352, Δ*htd2*+YEp352mtFam1-1, Δ*htd2*Δ*lip2*+YEp352, Δ*htd2*Δ*lip2*+YEp352mtFam1-1, Δ*htd2*Δ*gcv*3+YEp352 and Δ*htd2*Δ*gcv3*+YEp352mtFam1-1. Cells were grown for 16 hours, adjusted to an OD_600_ of 0.5 and spotted as 1x 1/10x, 1/100x and 1/1000x serial dilutions on a SCD-URA plate as a general growth control and on SCG-URA media supplemented with LA or C8. Plates were incubated at 30°C for 4 days. (B) Western blot analysis of extracts of mtFam1-1 complemented strains. Whole cell extracts were collected from cells expressing YEp352mtFam1-1 or YEp352 as a negative control, after 16 h of growth on YPG media at 30°C. Analysis was done by probing with anti-LA serum and anti-actin serum as a loading control. Whole cell extracts from wild type+YEp352, Δh*td2*+YEp352, Δ*htd2*+YEp352mtFam1-1, Δ*htd2*Δ*lip2*+YEp352 and Δ*htd2*Δ*lip2*+YEp352mtFam1-1 grown without supplements (−) or with 100μM C8 supplementation were analyzed with anti-LA serum. See Additional File Figure A1 for further data.

Growth of mtFAS-deficient Δ*htd2* cells expressing *FAM1-1* was barely detectable on unsupplemented SCG plates after a short 2-day incubation period (Figure 2A), but clearly visible after longer incubation times (Additional file Figure A12-14). In contrast, supplementation of SCG with either C8 or LA strongly improved growth (Figure 2A, Additional File Figure A1). Analysis of whole cell extracts by immunoblotting indicated restoration of protein lipoylation to wild type levels when C8 was added to the liquid broth (Figure 2B). YP-based media allowed residual growth of some mtFAS or lipoylation defective yeast strains on non-fermentable glycerol if fatty acid/LA activating enzymes were present in mitochondria. Also, we occasionally detected faint signals of lipoylated proteins in cell extracts even in the absence of C8 or LA supplementation, (Figure 2, Additional File Figures A1 – A3). This effect is likely due to the presence of small amounts of free C8 or LA in the rich YP-based media. Indeed, *the FAM1-1*-suppressed yeast strains show a graded improvement of growth in response to increasing concentrations of C8. (Additional File Figure A12-14). Taken together with previous results [27–29], these data are consistent with the hypothesis that free C8 is usually not available in yeast mitochondria. C8 and LA can be taken into the cells from the media, can enter mitochondria, and can be used for attachment to ketoacid dehydrogenase E2 subunits in mtFAS deficient yeast strains if a suitable activating enzyme is present.

### Fam1-1 suppression of mtFAS loss requires a functional lipoyl/octanoyl ligase Lip2

We wanted to further explore the rescue of Fam1-1-mediated respiratory growth and lipoylation of E2 targets in yeast mtFAS mutants. Toward this end, mutant strains combining the Δ*htd2* deletion with either Δ*lip2* or Δ*gcv3* were constructed and compared to the single mutant strains. Growth assays on supplemented media indicate that Fam1-1 only robustly rescues the respiratory growth of the Δ*htd2* single mutant strain (Figure 2A). The corresponding cell extracts analyzed by immunoblotting with anti-LA antiserum (Figure 2B) show that Fam1-1 expression in mitochondria did not restore lipoylation in any of the LA attachment pathway-deficient strains. Therefore, the data indicate that suppression of the respiratory growth defect by Fam1-1-activated C8 is a dependent on Lip2, the first enzyme in the LA synthesis/attachment process.

### Mitochondrially localized LplA supports respiratory growth and restores lipoylation in a mtFAS deletion strain supplemented with LA or C8

The *E. coli* lipoyl-protein ligase LplA catalyzes synthesis of lipoyl adenylate from free LA and ATP, and then bypasses the requirement for endogenous LA biosynthesis by transferring the cofactor to the apo-lipoyl-domains of 2-ketoacid dehydrogenases [30]. This enzyme has been reported to also accept C8 as a substrate, but with a 7-fold reduced specific activity compared to naturally occurring D-lipoate [30]. We tested whether the expression of mitochondrially targeted *E. coli* LplA could restore respiratory growth of the mtFAS deficient yeast Δ*htd2* strain (Figure 3A). Suppression of the Δ*htd2* strain respiratory growth defect by mitochondrial LplA, accompanied by recovery of lipoylation, was detected when the non-fermentable SCG medium was supplemented with LA or C8, but not without addition of these lipids. Rescue of growth was specific for LplA activity, since the negative control strain expressing an inactive LplA variant (LplAK133L) [11] showed no respiratory growth under any condition tested. The LA/C8 supplemented Δ*htd2*+LplA strain exhibits weaker respiratory growth compared to the WT control, presumably due to the lack of some other, as yet unidentified, mtFAS end product(s).

**Figure 3.**
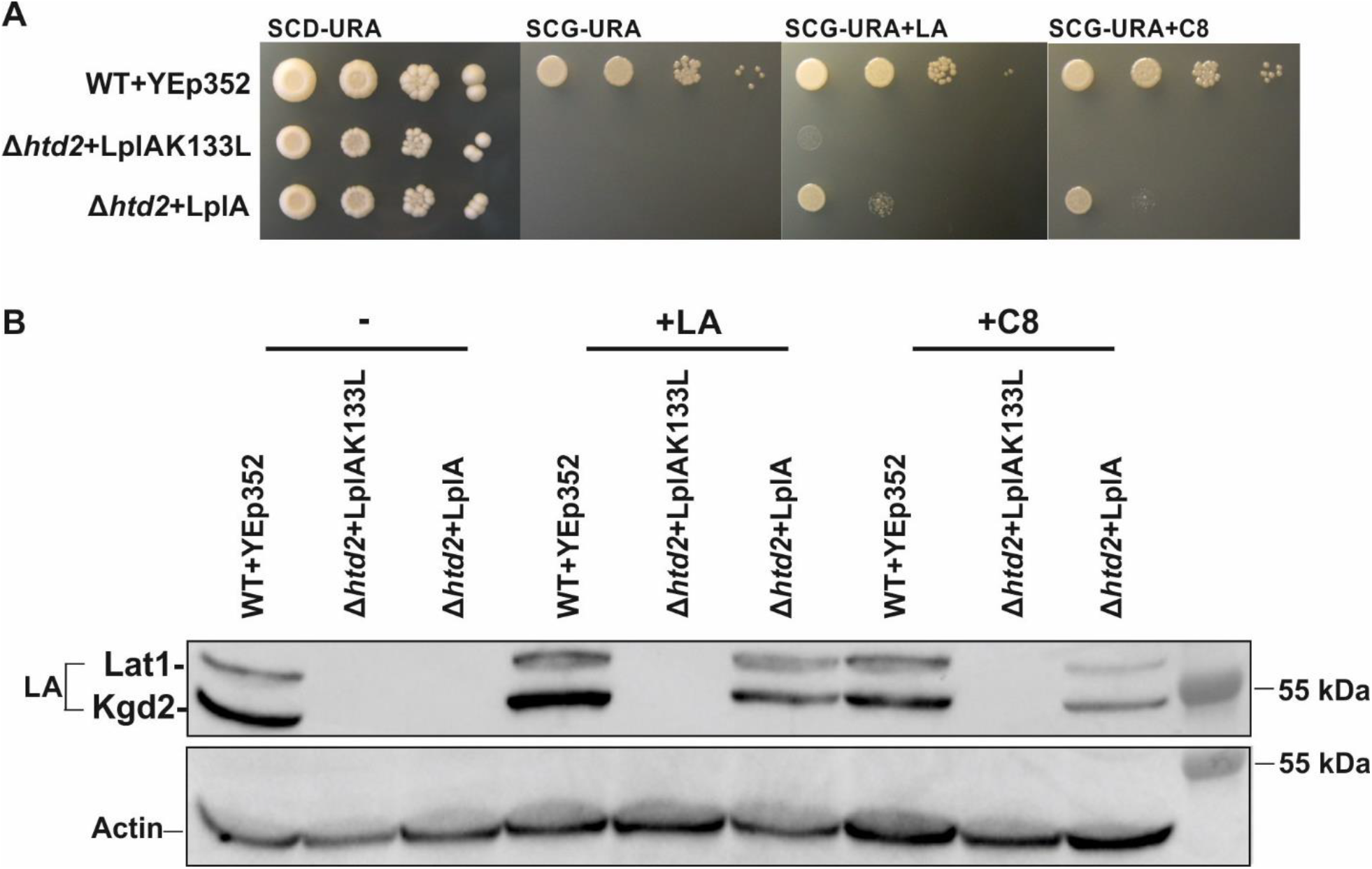
Analysis of the mtFAS-deficient Δ*htd*2 strain expressing mitochondrially targeted LplA. All plasmids transformed into the different strains are based on the YEp352 yeast episomal plasmid backbone. (A) Growth assays of the Δ*htd2* strain complemented with YEp352-LplA or YEp352-LplAK133L (negative control). Cells were grown for 16 hours, adjusted to an OD_600_ of 0.5 and spotted as 1x 1/10x, 1/100x and 1/1000x serial dilutions on SCD-URA media as a general growth control, and on SCG-URA media supplemented with LA or C8. Plates were incubated at 30°C for 4 days. (B) Western blot analysis of cell extracts from the same strains. Cells were grown for 16 h at 30°C on YPG, with and without supplements. Samples were probed with anti-LA serum using anti-actin as a loading control.

We surmised that rescue of Δ*htd2* strain growth by mitochondrially targeted LplA is dependent on the presence of exogenous LA and C8 in the media and is due to activation of these substrates by LplA for subsequent lipoylation of Lat1 and Kgd2. To test this hypothesis, we grew the Δ*htd2* strain expressing LplA on rich YPG media supplemented with either 100 μM LA or C8. Total cell protein was extracted and analyzed on immunoblots with anti-LA antiserum to assess protein lipoylation efficiency (Figure 3B). Consistent with the growth assay data, we were unable to detect lipoylation in the Δ*htd2*+LplA strain grown in unsupplemented medium. Although the growth of the cells expressing LplA in the presence of LA is not as robust as wild type (Figure 3A), the lipoylation of Lat1 and Kgd2 is almost fully restored. The increase in lipoylation was not quite as prominent with C8 supplementation, which may be explained by the lower activity of LplA utilizing C8 as a substrate [31]. No lipoylated proteins could be found in the negative control strain Δ*htd2*+LplAK133L expressing the inactive lipoyl-ligase variant. In line with the Fam1-1 suppression experiments, these data illustrate that free LA and C8 can traverse both the cell and mitochondrial membranes and, in the presence of the LplA enzyme, can be used for lipoylation of the PDH and KGD complexes.

### Mitochondrially localized LplA rescues growth and lipoylation in all lipoylation-deficient yeast mutants except the Δlip3 strain

We then investigated whether LplA was able to bypass one or more hierarchical biosynthetic steps of endogenous LA biosynthesis and protein lipoylation in yeast (Figure 1) [11] and rescue the phenotypes of lipoylation-deficient strains. The LplA-expressing Δ*lip5* strain was cultivated in the presence of LA, with a control culture grown in parallel on medium supplemented with C8. Respiratory growth of the Δ*lip5* strain was rescued by LplA only when the medium was supplemented with LA (Figure 4A, Figure S2). The immunoblot of the corresponding cell extracts with anti-LA serum showed restored lipoylation in the LA-supplemented sample (Figure 4B), indicating that the observed growth was the result of increased lipoylation.

**Figure 4.**
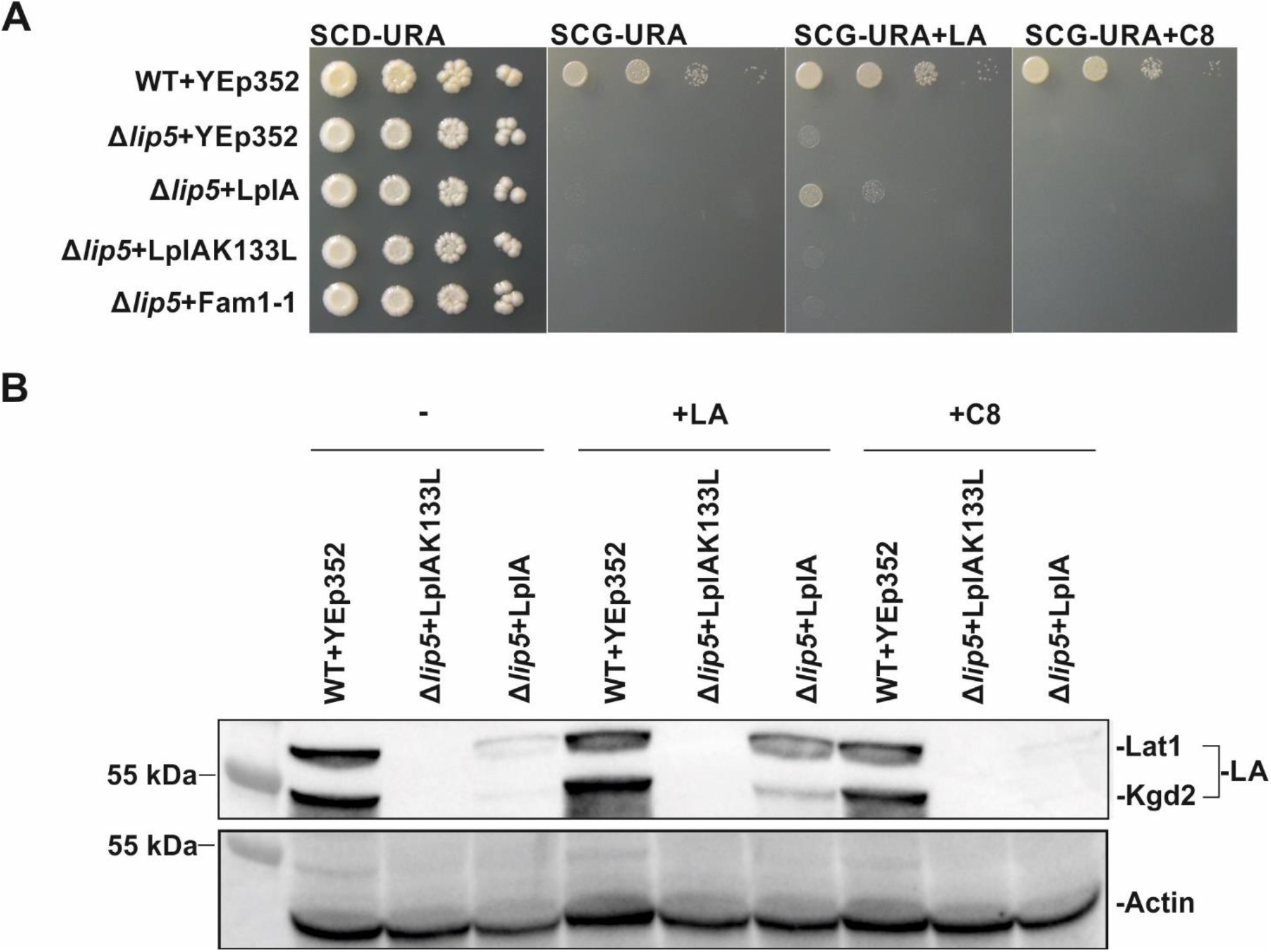
Analysis of the LA synthesis-deficient Δ*lip5* strain transformed with mitochondrially targeted LplA. All plasmids transformed into the different strains are based on the YEp352 yeast episomal plasmid backbone. (A) Growth assays of the Δ*lip5* strain complemented with YEp352-LplA or YEp352-LplAK133L (negative control). Cells were grown for 16 hours, adjusted to an OD_600_ of 0.5 and spotted as 1x 1/10x, 1/100x and 1/1000x serial dilutions on SCD-URA media as general growth control and on SCG-URA media supplemented with LA or C8. Plates were incubated at 30°C for 4 days. (B) Western blot analysis was carried out using cell extracts from the same strains. Cells were grown for 16 h at 30°C on YPG, with and without supplements. Samples were probed with anti-LA serum using anti-actin as a loading control.

Next we tested whether mitochondrial LplA would be able to restore growth and protein lipoylation in the other lipoylation defective yeast strains (Figure 5A, Figure S3A). Without LA or C8 supplementation, LplA partially reversed the respiratory growth defect of the Δ*lip2* strain. This weak growth was improved by the addition of LA or C8. Challenging immunoblots of protein extracts from the LplA-complemented Δ*lip2* strain with α-LA antiserum indicated that protein lipoylation was partially rescued by addition of C8, and fully rescued with LA supplementation (Figure 6A). No lipoylation could be detected in the samples grown in the absence of supplementation, (Figure 5A) (see Discussion).

**Figure 5.**
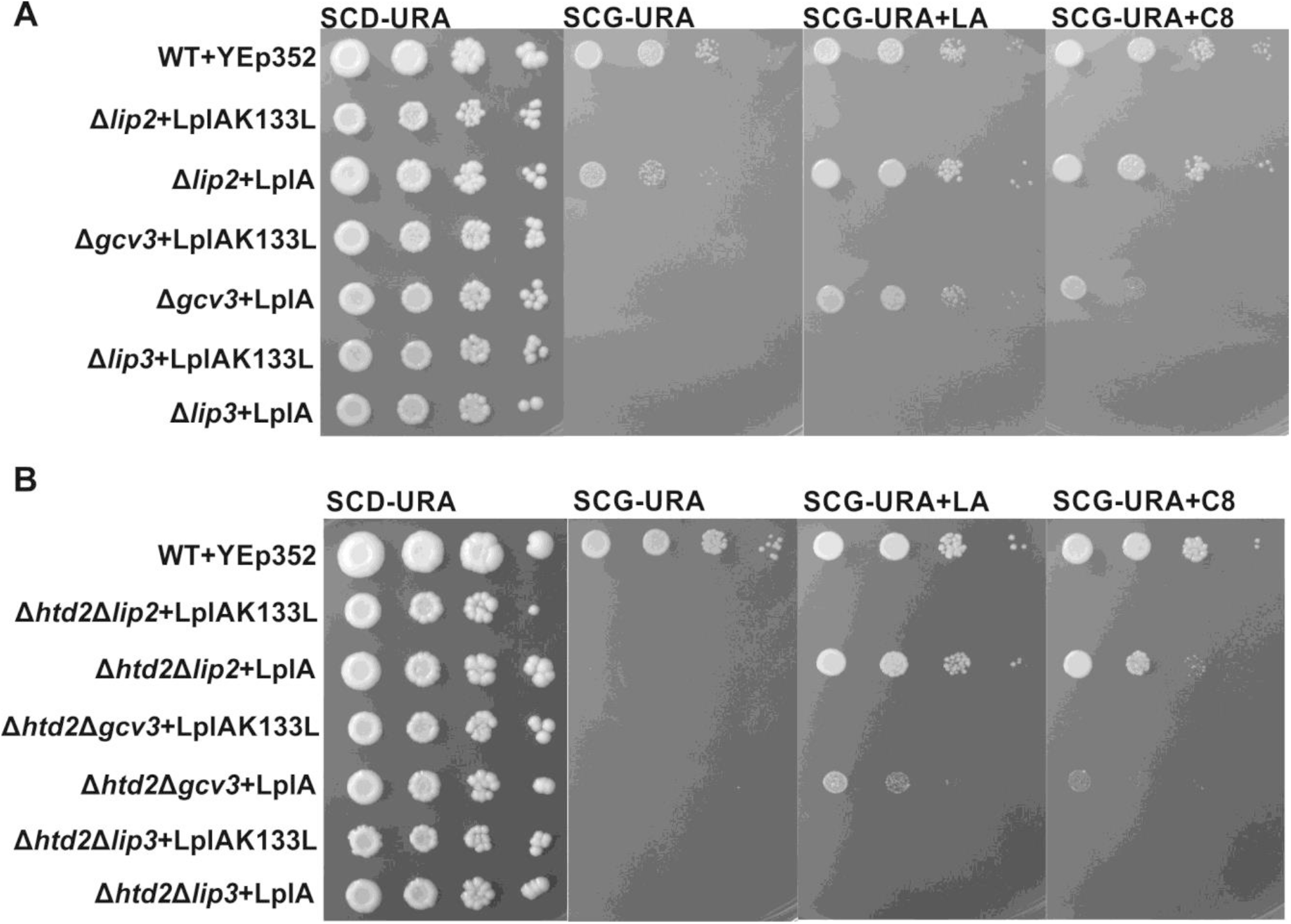
Growth assays of lipoylation deficient strains complemented with LplA. All plasmids transformed into the different strains are based on the YEp352 yeast episomal plasmid backbone. (A) Δ*lip2*, Δ*gcv3* and Δ*lip3* strains complemented with YEp352-LplA and YEp352-LplAK133L as a negative control. Cells were grown for 16 hours, adjusted to an OD_600_ of 0.5 and spotted as 1x 1/10x, 1/100x and 1/1000x serial dilutions on SCD-URA media as a general growth control, and on SCG-URA media supplemented with LA or C8. Plates were incubated at 30°C for 4 days. (B) Δ*htd2*Δ*lip2*, Δ*htd2*Δg*cv3* and Δ*htd2*Δ*lip3* complemented with YEp352-LplA, or YEp352-LplAK133L as a negative control, were spotted on a SCD-URA plate as a general growth control and on SCG-URA media with or without LA or C8. Plates were incubated at 30°C for 4 days.

**Figure 6.**
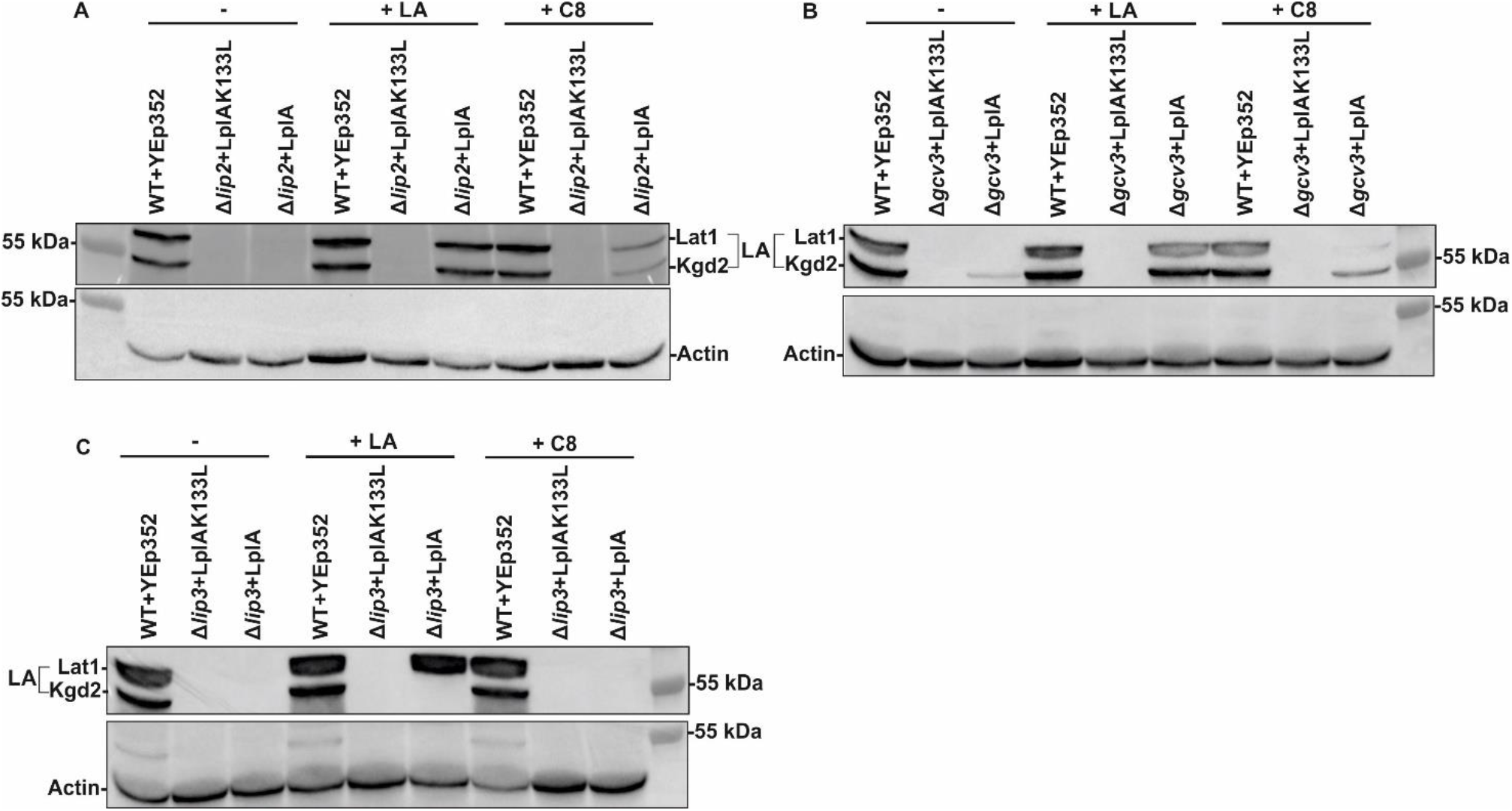
Western blot analysis of Δ*lip2*, Δ*gcv3* and Δ*lip3* yeast strains complemented with LplA. All plasmids transformed into the different strains are based on the YEp352 yeast episomal plasmid backbone. Whole cell extracts were collected from cells expressing either YEp352-LplA, or YEp352-LplAK133L as a negative control, after 16h of growth at 30°C on YPG media. Blots were analyzed with anti-LA serum, and anti-actin serum as a loading control. (A) Whole cell extracts from wild type+YEp352, Δ*lip2*+YEp352-LplA and Δ*lip2*+YEp352-LplAK133L grown without supplements (−), or with 100μM LA or 100μM C8, were analyzed with anti-LA serum. (B) Whole cell extracts from wild type+Yep352, Δ*gcv3*+YEp352-LplA and Δgcv3+YEp352-LplAK133L grown without supplements (−), with 100μM LA or 100μM C8 analyzed with anti-LA serum. (D) Whole cell extracts from wild type+Yep352, Δ*lip3*+YEp352-LplA and Δ*lip3*+YEp352-LplAK133L grown without supplements (−), or with 100μM LA or 100μM C8 were analyzed with anti-LA serum.

Schonauer *et al*. [11] previously showed that lipoylation of Lat1 and Kgd2 is abolished in the absence of Gcv3. We were curious whether rescue by LplA acted upstream or downstream of Gcv3. Respiratory growth of the Δ*gcv3* strain was restored by LplA when the cells were cultivated in the presence of LA, and to a lesser extent when grown with C8 (Figure 5A). Our immunoblot analysis revealed that Lat1 and Kgd2 are lipoylated in the Δ*gcv3* strain rescued by LplA and supplemented with LA or C8 (Figure 6B, Figure S3B). Thus, the physical presence of modified Gcv3 is not required for lipoylation under these conditions, and LplA must be acting downstream of Gcv3 to rescue lipoylation.

Surprisingly, in contrast to the Δ*lip2*, Δ*lip5* and Δ*gcv3* strains, LplA plus C8 or LA supplementation was not able to rescue the growth of the Δ*lip3* strain (Figure 5A). The reason for the non-complementation became apparent when we examined protein lipoylation in the LplA-expressing Δ*lip3* strain supplemented with LA or C8 by immunoblotting with anti-LA serum. We detected lipoylated Lat1, but not Kgd2 (Figure 6C). This result implies that *E. coli* LplA does not recognize, or does not have access to, the lipoylation domain of Kgd2, impeding the rescue of respiratory deficient growth of the Δ*lip3* strain. Thus, LplA is not able to lipoylate both E2 target proteins in yeast.

Our observation of partial rescue of Δ*lip2* strain respiratory growth in the absence of supplementation or by scavenging C8 or LA from the medium, prompted us to test whether LplA can use acyl-groups produced by the mtFAS pathway by deleting *HTD2*. The growth assays show that in the Δ*htd2*Δ*lip2* background, LplA no longer rescues the respiratory defect of Δ*lip2* when grown without supplements (SCG-URA, Figure 5B). Thus, the rescue of respiratory growth of the Δ*lip2* strain by LplA in the absence of supplements is mtFAS-dependent.

### The lipoyl domain of Lat1 can substitute for Gcv3 as a donor for lipoylation of Kgd2 by Lip3

If LplA can only recognize Lat1 and not Kgd2, how is Kgd2 lipoylated in Δ*lip2*, Δ*lip5,* and especially Δ*gcv3* strains? Can the wild type Lip3 in these deletion strains work by transferring LA directly from LplA to Kgd2? Or, perhaps, is Lip3 scavenging LA from lipoylated Lat1? We decided to test these hypotheses directly.

To eliminate the possibility of transfer from Lat1, thus testing the alternative hypothesis that Lip3 can accept activated LA directly from LplA, we generated the Δ*gcv3*Δ*lat1* double deletion strain and expressed mitochondrial LplA in this strain. Interestingly, the growth defect of this strain on SCG media was rescued by LA or C8 supplementation (Figure 7A), but absolutely no lipoylation was detected (Figure 7B). Thus, Lip3 is unable to use LplA-activated LA directly for lipoylation of Kgd2. Therefore, we hypothesized that the restoration of lipoylation by LplA we observed in the LA-supplemented Δ*gcv3* strain (Figure 6B) was due to a previously uncharacterized ability of Lip3 to transfer LA from Lat1 to Kgd2. To test this idea, we constructed a strain expressing Lat1 with the C-terminal residues 185-482 deleted. The remaining 184 N-terminal residues comprise the lipoyl domain, followed by a flexible linker region and the E1-binding domain. This construct lacks the large E2 C-terminal catalytic domain. Lat1Δ185-482 was integrated into the genome of the Δ*gcv3*Δ*lat1* strain and we tested whether LplA plus supplementation of the medium with LA would restore Kgd2 lipoylation in this strain. Immunoblot analysis revealed the presence of lipoylated Kgd2 when the medium was supplemented with LA (Figure 7C), which implies that Lip3 can function as a lipoamide transferase, using the Lat1 lipoyl domain construct as a substrate for the lipoylation of Kgd2 by Lip3. Expression of truncated Lat1 in the presence of LplA and LA also rescued growth, even though PDH is not active in these cells. In the absence of PDH activity, mitochondrial acetyl-CoA is provided to the intact TCA cycle by the pyruvate dehydrogenase bypass pathway: sequential action of pyruvate decarboxylase, acetaldehyde dehydrogenase and acetyl-CoA synthase, and carnitine acetyltransferase [32].

**Figure 7.**
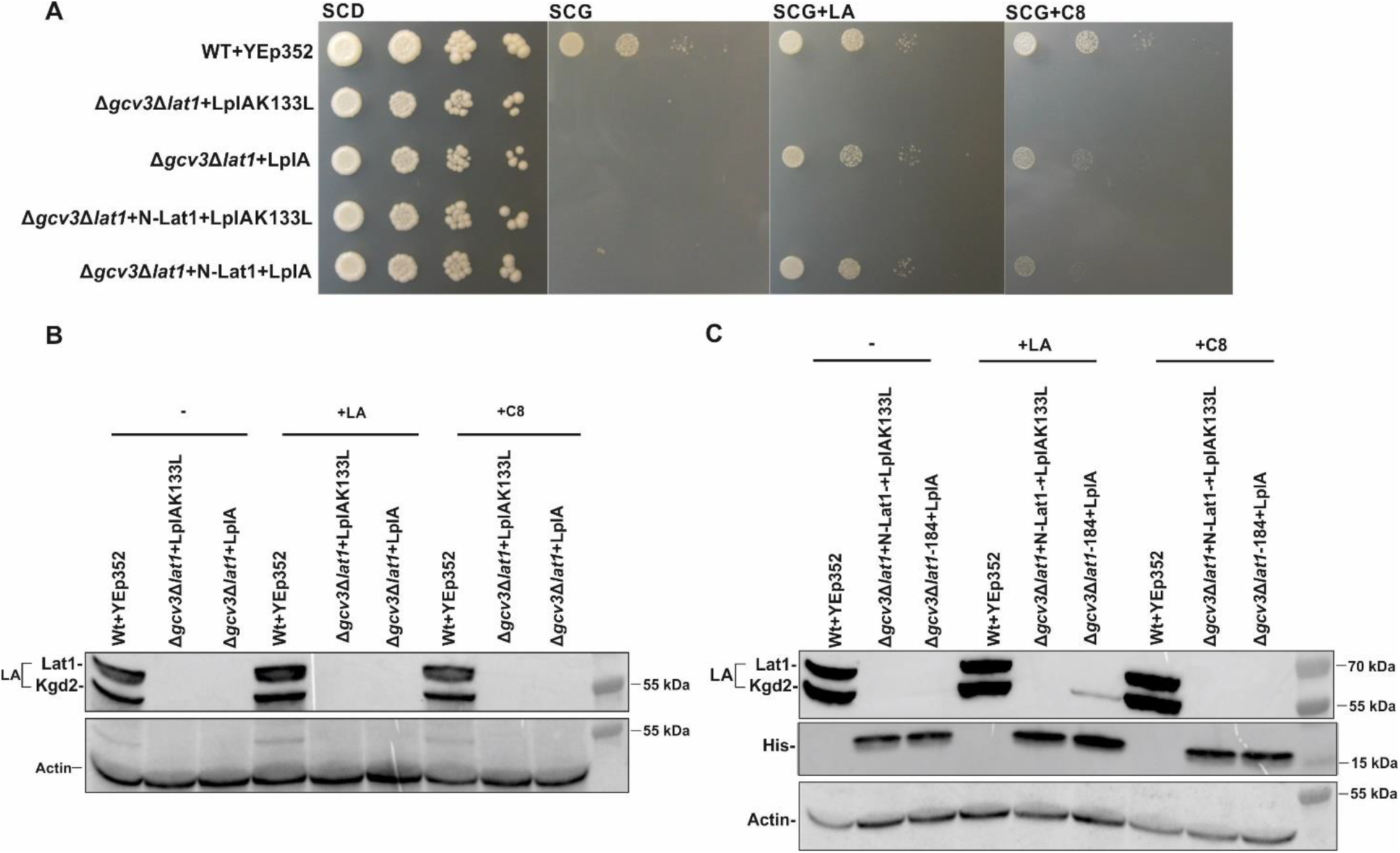
Growth assay and western blot analysis of Lip3 transfer activity with truncated Lat1 as the lipoyl donor. All plasmids transformed into the different strains are based on the YEp352 yeast episomal plasmid backbone. (A) Growth assay of wild type+YEp352, Δ*gcv3*Δ*lat1*+YEp352-LplAK133L, Δ*gcv3*Δ*lat1*+YEp352-LplA, Δ*gcv3*Δ*lat1 +* N-Lat1 + YEp352-LplAK133L and Δ*gcv3*Δ*lat1* + N-Lat1+YEp352-LplA. Cells were grown for 16 hours, adjusted to an OD_600_ of 0.5 and spotted as 1x 1/10x, 1/100x and 1/1000x serial dilutions SCD-URA as the general growth control, as well as on SCG-URA, SCG-URA+100 μM LA or SCG-URA+100 μM C8. Plates were incubated for 4 days at 30°C. For western blot analysis, whole cell extracts were collected from cells grown in YPG with and without 100 μM LA or C8 supplementation. (B) Whole cell extracts of wild type+YEp352, Δ*gcv3*Δ*lat1*+LplA and Δ*gcv3*Δ*lat1*+LplAK133L strains were analyzed with anti-LA serum and anti-actin serum for the loading control. Please note that the white line separating lane 4 and 5 in the lipoylation signal data is not a splicing artefact, but an artefact of the overlay with the image file containing the size marker information. All samples are from the same western blot and are continuous (see Additional File Figure A4) and the identical membrane was used to probe for the actin loading control. (C) Whole cell extracts of the Δ*gcv3*Δ*lat1* strain expressing the His6- and C-protein double-tagged N-Lat1 lipoyl domain plus either the YEp352-LplA or YEp352-LplAK133L plasmids were analyzed on immunoblots. The anti-His6 signal indicates the presence of the His-tagged N-Lat1 lipoyl domain protein fragment. The blots were reacted with anti-LA serum, anti-His serum, and anti-actin serum for the loading control.

### The yeast lip2 and lip3 mutations are complemented by the human homologs LIPT2 and LIPT1 respectively

Because the pathway for protein lipoylation is highly conserved, and the homologs for the yeast enzymes have been identified in humans [7, 12, 13, 20], we tested whether the human enzymes could complement the respective yeast mutant strains. The *LIPT2* and *LIPT1* genes were cloned into a yeast expression vector [33] and the resulting plasmids were transformed into yeast deletion strains as follows: YEp352-*LIPT2* into Δ*lip2* and YEp352-*LIPT1* into Δ*lip3*. Growth assays on plates show that the human enzymes complement the respiratory growth defects of the yeast deletion strains (Figure 8A, B).

**Figure 8.**
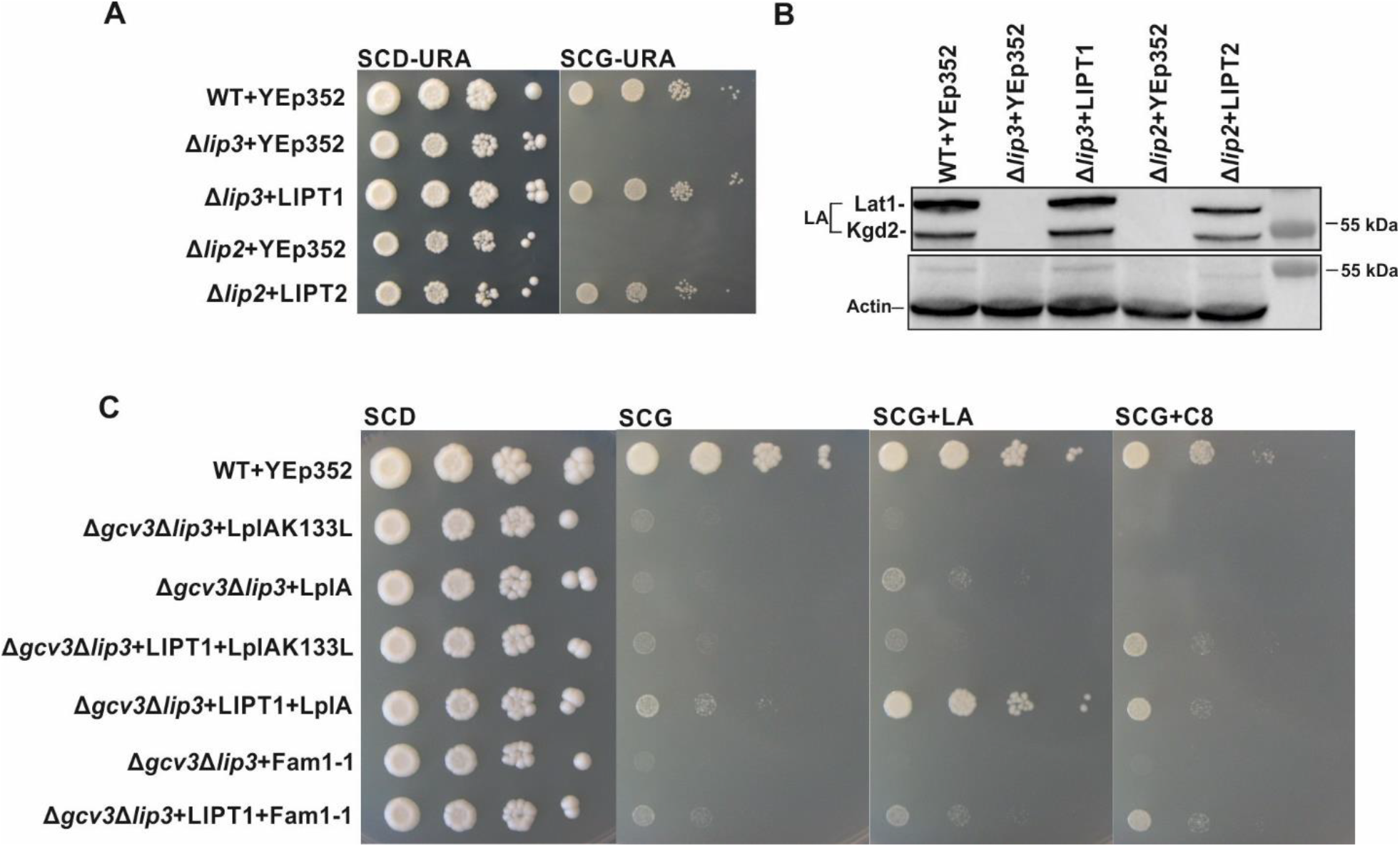
Complementation of Δ*lip2*and Δ*lip3* yeast strains by human LIPT1 and LIPT2. All plasmids transformed into the different strains are based on the YEp352 yeast episomal plasmid backbone. (A) Growth assays: wild type+YEp352, Δ*lip3*+YEp352, Δ*lip3*+YEp352hsLIPT1, Δ*lip2*+YEp352 and Δ*lip2*+YEp352hsLIPT2. Cells were grown for 16 hours, adjusted to an OD_600_ of 0.5 and spotted as 1x 1/10x, 1/100x and 1/1000x serial dilutions on SCD-URA media as a general growth control, and on SCG-URA media. Plates were incubated at 30°C for 4 days. (B) Western blot analysis of cell extracts of wild type+YEp352, Δ*lip3*+YEp352, Δ*lip3*+YEp352HsLIPT1, Δ*lip2*+YEp352 and Δ*lip2*+YEp352HsLIPT2 with anti-LA serum. (C) Growth assays of wild type, Δ*gcv3*Δl*ip3*+YEp352-LplAK133L, Δ*gcv3*Δ*lip3*+YEp352-LplA, Δ*gcv3*Δ*lip3*+YEp352-LIPT1+YCp22-LplAK133L, Δ*gcv3*Δ*lip3*+YEp352-LIPT1+YCp22-LplA. Cells were spotted on a SCD-URA-TRP media as a general growth control, and on SCG-URA-TRP media without supplements or supplemented with 100 μM LA or 100 μM C8. Plates were incubated at 30°C for 4 days.

**Figure 9.**
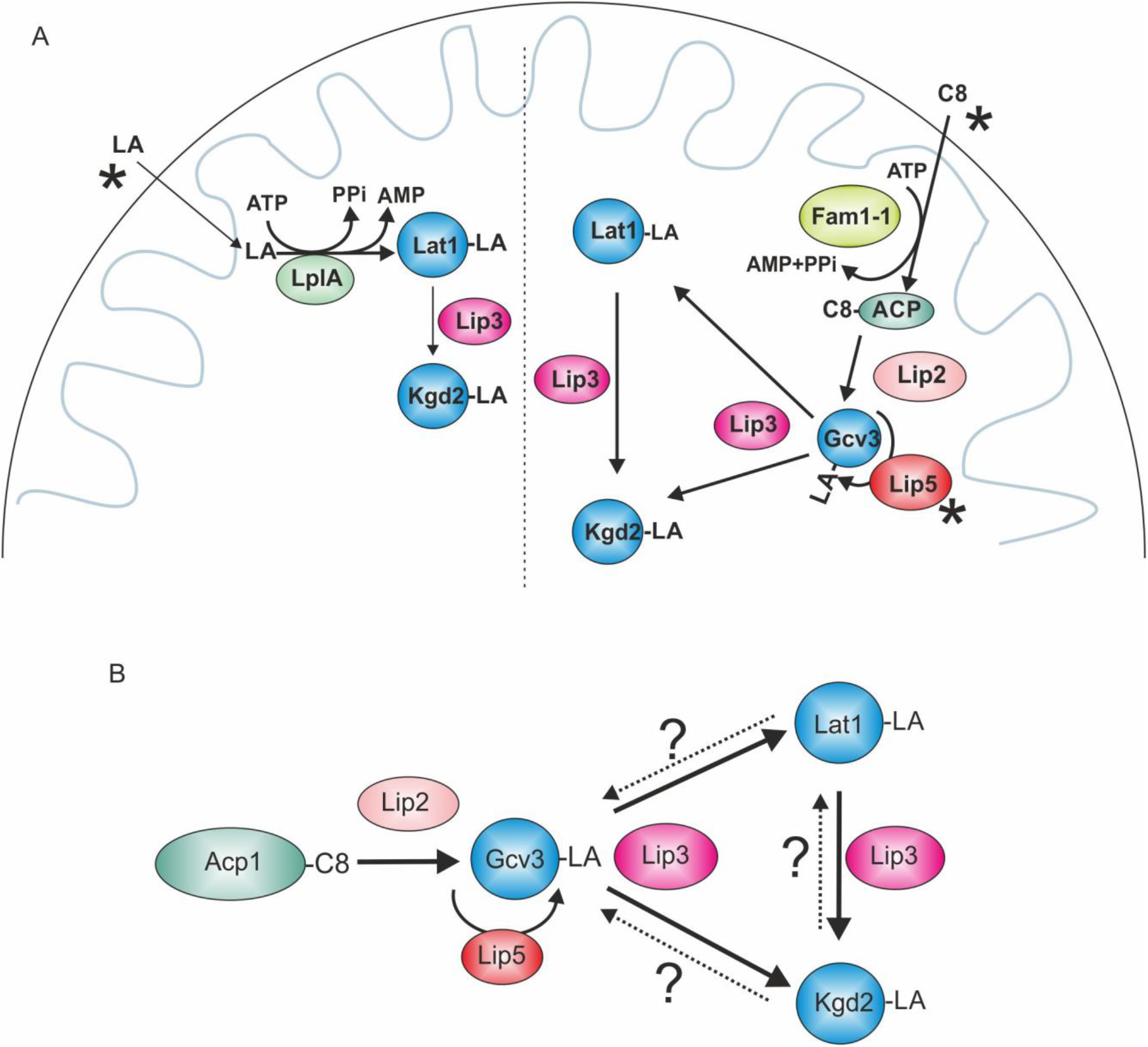
Models for lipoylation mechanisms. A) Lipoylation in yeast strains expressing suppressor constructs (mitochondrially targeted LplA or Fam1/Faa2) under supplementation conditions. Left: After LA has passed through the mitochondrial membranes, LplA activates it with AMP and transfers the lipoyl moiety onto Lat1, which is used as a substrate by Lip3 for Kgd2-lipoylation. Right: After entering mitochondria, C8 is activated by Fam1-1, possibly by direct attachment to ACP. Octanoyl-ACP serves as a substrate for Lip2 and the C8 is transferred onto Gcv3. Lip5 and Lip3 use octanoyl-Gcv3 for attachment of thiol groups and as a substrate for transfer to the PDH and KGD-complexes, respectively. B) Lipoylation events under native conditions. Lip2 transfers C8 from Acp1 to Gcv3 and Lip5 adds the thiol groups to carbons 6 and 8, forming LA. Lip3 transfers the LA moiety onto Lat1 and Kgd2. The novelty of this model, based on the data presented here, is that Lip3 can also transfer LA from Lat1 to Kgd2. We postulate that this may serve as an additional layer of regulation to match the prevailing energy flux in the cell with PDH and KGD-complex activities. This activity of Lip3 could serve to quickly fine-tune the relative activities of PDH and KGD by simple transfer of the lipoyl moiety from Lat1 to Kgd2, for example in glucose-deprived conditions where the need for PDH-complex activity is reduced.

### The yeast Δlip3 strain expressing human LIPT1 and E. coli LplA responds to LA and C8 supplementation

The plasmid-borne copy of LIPT1 complements the yeast Δ*lip3* strain and thus we were curious to determine whether the human lipoyl-protein ligase also responded to LplA activated LA/C8. To this end, we transformed the double deletion strain Δ*gcv3*Δ*lip3* with theYEp352-LIPT1 plasmid in combination with a plasmid carrying either LplA or LplAK133L. The growth assays show that the respiratory growth defect of the Δ*gcv3*Δ*lip3* strain can be rescued by the expression of LIPT1 and LplA (Figure 8C). Growth rescue is partial in the absence of supplementation and is clearly improved by the addition of LA to the growth media.

## DISCUSSION

In light of the emergence of disease-causing inborn errors in humans associated with lipoylation defects [7, 12, 16, 20], there is a pressing need to understand the underlying principles of LA/C8 transfer to key mitochondrial enzymes and the susceptibility of the lipoylation process to substrate supplementation. The existence of an LA-scavenging pathway in eukaryotes has been controversial, but evidence from several studies in yeast are in agreement that supplementation of the medium with free fatty acid or LA does not provide growth enhancement [6, 9]. The ability of higher eukaryotes to bypass the need for mtFAS-synthesized LA is even more controversial than in yeast. Similar to the yeast models, several studies with human cell lines or mouse models with deficiencies in LA synthesis [34, 35], Fe-S cluster metabolism [20, 36–38] or protein lipoylation [12, 13, 20], do not support the existence of a pathway enabling mammalian cell use of exogenous LA for lipoylation of mitochondrial enzyme complexes. A single study did report that LA supplementation partially restored mitochondrial dysfunction in HEK293 cells that had been treated with the C75 inhibitor of the mtFAS-condensing enzyme OXSM, but data demonstrating restoration of protein lipoylation levels were not provided [39]. In agreement with most of the studies mentioned above, our experiments showed that yeast strains deficient in each step of the lipoylation pathway did not regain significant respiratory growth upon addition of either LA or C8 to the growth medium, and none of the target proteins exhibited an increase in lipoylation (Fig.2, Fig. 3).

It was clear from the collective data that free fatty acids either do not enter yeast cells and mitochondria, or they are able to enter but they are not activated by covalent attachment to AMP, CoA or ACP for presentation as a substrate for the lipoylation pathway. To test the hypothesis that the deficiency was in activation, we employed the *FAM1-1* suppressor allele or *E. coli* LplA endowed with a yeast mitochondrial targeting signal and expressed in *S. cerevisiae* strains with deletions in each of the genes encoding successive lipoylation pathway components. *FAM1-1* activates fatty acids by attachment to CoA or to ACP [40]. The dependence of C8 supplementation on Lip2 (required for transfer of C8 to Gcv3) is consistent with this conclusion. Intriguingly, LA supplementation also enhances growth of mtFAS deficient strains on non-fermentable carbon sources (Fig. 2), in a Lip2-dependent manner. This result is consistent with initial attachment of LA by Fam1-1 to ACP, and subsequent transfer to Gcv3. The “business end” carboxyl group of LA and C8, used in the thioester bond to ACP or the amide bond to Gcv3 or E2 subunits of PDH of KGD, is identical for both molecules. Therefore, observing LA functioning as a substrate for Fam1-1 is not surprising. Likewise, LplA activates free lipoic acid by covalent attachment to AMP [41], and is capable also of activation and transfer of C8 [31]. When localized to yeast mitochondria, LplA rescued the respiratory deficient growth phenotype of the Δ*lip2* strain even without LA or C8 supplementation, but detectable lipoylation of the E2-subunits was absent under these experimental conditions. Thus, LplA bypassed the need for Lip2 for respiratory growth without addition of supplements, but restoration of protein lipoylation was only observed with LA or C8 supplementation. Unlike Lip2, the need for Gcv3 was bypassed by LplA only in the presence of added LA or C8. That LplA can rescue the respiratory growth-deficient phenotypes of both the Δ*lip2* and Δ*gcv3* strains reveals that Lip3 substrate usage is not restricted to lipoylated Gcv3 (the yeast GCS H protein) as the LA donor, implying that both LA and C8 must be activated by AMP and available for attachment to E2 subunits in this form.

Unexpectedly, LplA did not rescue growth of the Δ*lip3* strain, even though Lip3 and LplA share substantial sequence similarity [11]. Close examination of the immunoblots suggested that LplA is capable of lipoylating only the yeast Lat1 lipoyl domain, and not the Kgd2 lipoyl domain. In combination with our finding that lipoylation in a Δ*gcv3*Δ*lat1* strain was not restored by LplA, and underscored by our observation that lipoylated Kgd2 was detected in the LplA-complemented Δ*gcv3*Δ*lat1* strain expressing a His-tagged, truncated Lat1 lipoyl-domain, these results are consistent with the ability of Lip3 to efficiently transfer LA from Lat1 to Kgd2 *in vivo* (Figure 7).

Our complementation assays also indicate that the LA synthase Lip5 can recognize C8 attached to proteins other than Gcv3. In our growth assays, the only lipoylation deficient strain background that was not rescued by C8 supplementation in the presence of activating enzyme was *Δlip5*. This indicates that in *LIP5* strains, including *Δgcv3*, C8 is a substrate for Lip5-mediated sulfur addition and conversion to LA after attachment to, for example, Lat1. However, LA supplementation consistently leads to a more pronounced restoration of lipoylation in comparison to C8 supplementation. This observation may be partly explained by the seven-fold lower specific activity of LplA using C8 as a substrate in comparison to LA [30].

Our results are in agreement with the model that Lip2 specifically modifies only the yeast H-protein (Gcv3) during the protein lipoylation process [11], although it can modify the bacterial lipoylated complexes when expressed in LipB and LplA deficient *E. coli* strains [42]. Results from patient studies indicate that the human octanoyl transferase LIPT2 functions similarly to yeast Lip2. Fibroblasts obtained from patients suffering from mutations in *LIPT2* show strongly reduced lipoylation of the 2-oxoacid dehydrogenase complexes [20]. The LIPT2-deficient patients also showed elevated blood plasma glycine levels, which is indicative of defective glycine cleavage system activity and implies that the LIPT2 mutations also impede GCS-function, as would be expected based on the yeast data. In contrast to LIPT2-deficient patients, individuals suffering from defects in the lipoyltranferase LIPT1 (the Lip3 homolog) do not show elevated plasma glycine levels and the lipoylated H-protein of GCS can be detected in immunoblots of protein extracts from fibroblasts of these patients [12, 13]. In addition, the levels of PDH- and KGD-complex lipoylation are severely reduced in LIPT1 deficiency patients, indicative of conservation of lipoylation mechanisms between humans and yeast, in which GCS H protein lipoylation is a prerequisite for modification of other E2 subunits. Our work presented here reaffirms a role for the lipoylated H protein as a substrate for LA transfer rather than as a structural component of the lipoylation machinery, since the deletion of *GCV3* does not interfere with transfer of LA from activated C8/LA in LplA-expressing yeast cells.

Throughout our experiments, the level of lipoylation detected by LA antiserum is not always proportional to the level of growth rescue on respiratory media, in that modest growth improvement is a more sensitive indicator of low levels of lipoylation. Acyl-ACPs generated by mtFAS are required for respiratory growth of yeast cells via their contribution to mitochondrial Fe/S-cluster synthesis, mitochondrial protein synthesis, and assembly of respiratory chain complexes [43]. However, growth of the *Δlip2*+LplA strain and the Δ*gcv3*Δ*lat*1+LplA strain on glycerol without supplements (and displaying no detectable lipoylation) (Figure 7) calls for an additional and/or alternative explanation. Perhaps LplA mobilizes a mtFAS product for RNase P modification as suggested by Schonauer et al. [24], or another mitochondrial target.

In terms of utilization of a mtFAS derived acyl substrate, it is unlikely that LplA would use acyl-ACP as a direct substrate. However, a functional mtFAS is required for growth of the Δ*gcv3* and the *Δlip2* yeast strains expressing LplA on SCG media. This observation calls for the presence of an enzyme in mitochondria capable of releasing acyl moieties from ACP. The freed acyl groups would then be both activated and transferred by LplA, allowing full rescue of growth, even in the absence of supplements. Such a scenario would explain why, in the background of an additional mtFAS deficiency (Δ*htd2Δlip2*) where no acyl-ACP is available, the rescue of the *lip2* or *gcv3* deficiency by LplA is dependent on LA or C8 supplementation (Figure 5B).

Very weak hints of growth may be detected for some mutant strain backgrounds when minimal medium is supplemented with LA (*e.g.* Figure 2), an effect that is enhanced on YP-based media (Additional File Figure A1). Although we are not inclined to score these yeast “ghost” imprints as positive for growth, we would like to point out that *FAM1/FAA2* was recently identified as one of many yeast genes possibly encoding a mitochondrially localized variant of a protein, produced by translation initiation from a non-canonical start codon [44, 45]. Mitochondrial localization of minute amounts of native Fam1/Faa2, due to altered translation initiation under certain environmental or metabolic conditions, may allow slight growth, though we have yet to explore this possibility. As mtFAS/LA synthesis pathways appear to be organized in positive feedback loops, mitochondrially localized Faa2 might act as a failsafe that prevents a catastrophic system breakdown when mitochondrial acetyl-CoA levels are diminished.

In contrast to lackluster growth in response to supplementation in previous studies, and no restoration of lipoylation, expressing a mitochondrially-localized activating enzyme, such as the fungal fatty acyl-CoA ligase Fam1-1, or the bacterial LA ligase LplA, made C8 and LA available as substrates for protein lipoylation and improved growth. These data also imply that both compounds can traverse the mitochondrial membranes, but yeast mitochondria normally lack enzymes which make them available to the LA-attachment machinery.

Our new results also have a few surprising implications concerning *in vivo* substrate usage of LplA, Lip5 and Fam1-1. Our data suggest that LplA can use C8 as a substrate, and that Lip5 is able to modify the octanoyl moiety to the lipoyl form not only on Gcv3, but also on the E2 subunits Lat1 and Kgd2.

Taken together, we demonstrate that LA and C8 enter mitochondria, where they are used as substrates for protein lipoylation in the presence of an appropriate activating enzyme. In addition to the reported activity of transferring LA from Gcv3/GCS H protein to Lat1 and Kgd2, we show that Lip3 can operate as a transamidase capable of transferring LA from the PDH complex to the KGD complex. This result may indicate a regulatory role for Lip3 in adjusting the relative activities of the two complexes to match the metabolic status of the cell. In this model, Lip3 can be dynamic in fine tuning the lipoylation levels between the PDH- and KGD-complexes [46]. In addition, our data indicate that the eukaryotic lipoyl synthase Lip5 must be capable of thiolation of C8 not just on Gcv3, but also directly on the E2 subunits of the other ketoacid reductase complexes. We show that the human LIPT2 and LIPT1 are active in yeast and can replace the function of the homologous yeast enzymes in protein lipoylation, Lip2 and Lip3 respectively. LIPT1 also responds positively to LA supplementation in the presence of LplA by restoring respiratory growth in a yeast *gcv3* deletion background. When lipoyl transferase/transamidase is present in mitochondria, mtFAS/lipoylation deficient eukaryotic cells respond positively to artificially activated C8/LA substrates. We suggest that our experimental setup in yeast is a robust and simple platform to study the human enzymes and different substrates they can accept *in vivo* for protein lipoylation. It may also be developed into a tool for the identification of potential drugs for patients suffering from protein lipoylation deficiencies, while maintaining an alternative functional octanoyl/lipoyl transferase system.

## CONCLUSIONS

Using yeast as a model organism, we demonstrate that the eukaryotic lipoylation pathway can use free LA and C8 as substrates when fatty/lipoic acid activating enzymes are targeted to mitochondria. Our data also indicate that Lip3 transferase recognizes lipoylated Lat1, the E2 subunit of PDH, as a substrate for LA transfer to Kgd2, the E2 subunit of KGD, in addition to recognizing the major substrate Gcv3-LA. The mammalian homologs LIPT1 and LIPT2 fully complement the corresponding yeast deletion mutations, and respond similarly to activation enzyme and substrate supplementation, which suggests conservation of the eukaryotic lipoylation pathway from yeast to humans. These findings suggest it may be possible to develop small molecule therapeutics that are accepted by lipoyl transferases and prove effective in the treatment of LA or mtFAS deficiency-related disorders.

## METHODS

### Strains, media and growth conditions

For strains used in this study see Table 1. All strains are isogenic and based on strain W1536 5B [27]. The wild type yeast strain carried URA plasmid YEp352, an episomal, multicopy yeast plasmid [47]. For supplementation studies, yeast cells were grown either on rich or synthetic media. Rich media were all YP (1% yeast extract, 2% peptone) plus either 2% glucose for YPD or 3% glycerol for YPG. Synthetic complete media contained 2% glucose for SCD, or 3% glycerol for SCG. SCD and SCG media lacking the appropriate amino acids were used for plasmid selection. YP-based media appeared to contain small amounts of C8 or lipoic acid, thus growth assays on synthetic media provided clearer data for strain comparison. YPG and SCG were supplemented with 100 μM LA or C8 from a stock prepared in 70% ethanol. When needed during strain construction, media were supplemented with 300 μg/ml geneticin or 250 μg/ml of hygromycin B. Solid media contained 2% agar. All liquid cultures were grown at 30°C with vigorous shaking. The lipoylated proteins were much more easily detected in extracts of cells grown on glycerol than on other carbon sources. In order to be able to include also the negative control strains, the last growth step for the analysis of lipoylation state was carried out on YP-based media. This strategy also allowed the acquisition of cells from unsuppressed/uncomplemented knockout strains, which show weak growth on complex media.

**Table 1.** Strains used in this study (isogenic, based on W1536 5B)

| 516 | *S. cerevisiae* | Genotype | Reference |
| --- | --- | --- | --- |
| 517 | W1536 5B | *MAT a, Δade2, Δade3, can1-100, his3-11, 15; leu2-3, 112; trp1-1, ura3-1* | [27] |
| 518 | *Δhtd2* | *yhr067w::kanMX4* | [29] |
| 519 | *Δlip2* | *ylr239c::kanMX4* | [29] |
| 520 | *Δgcv3* | *yal044c::kanMX4* | [29] |
| 521 | *Δlip3* | *yjl046w::kanMX4* | [29] |
| 522 | *Δlip5* | *yor196c::kanMX4* | This study |
| 523 | *Δhtd2Δlip2* | *yhr067w::kanMX4, ylr239c::kanMX4* | This study |
| 524 | *Δhtd2Δgcv3* | *yhr067w::kanMX4, yal044c::kanMX4* | This study |
| 525 | *Δhtd2Δlip3* | *yhr067w::kanMX4, yjl046w::kanMX4* | This study |
| 526 | *Δgcv3Δlat1* | *yal044c::kanMX4, ynl071w::HPHMX* | This study |
| 527 | *Δgcv3Δlat1+N-Lat* | *yal044c::kanMX4, ynl071w::HPHMX, N-Lat integrated* | This study |
| 528 | *Δlip2Δlip3* | *ylr239c::kanMX4, yjl046w::kanMX4* | This study |
| 529 | *Δgcv3Δlip3* | *yal044c::kanMX4, yjl046w::kanMX4* | This study |

YEp352-Fam1-1, which encodes a mitochondrially localized Faa2 fatty acyl-CoA ligase from the yeast *CTA1* promoter, was described previously in Kursu et al. 2013 [29]. YEp352-LplA and-LplAK133L were described previously [48].

Manipulation of DNA and plasmid constructions were carried out using standard techniques. All new plasmid constructs were verified by sequencing. The cDNA clones for human LIPT1 and LIPT2 were purchased from Creative Biogene. The plasmids YEp352-LIPT1 and YEp352-LIP2 were constructed by cloning the coding sequences of LIPT1 and LIPT2 into the YEp352-CTA1 plasmid, after prior removal of the *CTA1* ORF with SphI/XbaI and HindIII. LIPT1 and LIPT2, digested with the same enzymes, were ligated into the vector, which retained the *CTA1* promoter and terminator sequences. No additional mitochondrial targeting sequences were introduced into the constructs.

Yeast integrative vector YIp349-LAT1-ST7 was used as a backbone for cloning and integrating a sequence encoding the N-terminal 184 amino acids of Lat1 into the Δ*gcv3* strain. The vector YIp349-LAT1-ST7 was constructed in collaboration with Professor Alexander Tzagoloff, containing the sequence encoding full length Lat1 followed by the CH-tag (C-reactive peptide and 6 x His residues) [49]. For constructing YIp349-N-Lat1-CH, the YIp349-LAT1-ST7 plasmid was used as a template for PCR-amplification of the 400 bp upstream untranslated region of *LAT1* followed by base pairs 1-552 of the coding sequence. YIp349-LAT1-ST7 was digested with SacI and PstI to remove the full-length *LAT1*. The PCR product was digested with SacI and PstI and ligated upstream and in-frame with the CH-tag.

Plasmids YCp22-LplA and YCp22-LplAK133L were constructed by digesting plasmids YCplac22 and YEp352-LplA/LplAK133L with EcoRI, purifying the digested fragments and ligating LplA/LplAK133L into the YCp22-vector. Fam1-1 was ligated into YCp22 in a similar manner.

All results shown are representative data, based on at least three independent biological repeats, with exception of the Δ*gcv3*, Δ*lat1* experiments shown in Figure 7, which were only done twice (Additional Files Figures A5 – A15).

### Disruption of open reading frames

Each open reading frame was deleted by homologous recombination with the *kanMX4* cassettes amplified from the respective deletion strains of the EUROSCARF collection as described by Kursu et al. [29] or amplification of the *hphMX* from plasmid pAG32 (RRID: Addgene_35122) with oligonucleotides introducing homology to target genes on the 5’ and 3’ ends of the PCR product.

Generation of strain W1536 5B Δ*gcv3*Δ*lat1* and W1536 5B Δ*gcv3*Δ*lat1* + N-Lat1.

The HPHMX gene (hygromycin resistance) was amplified from a plasmid pAG32 with primers containing 5’ and 3’ UTR sequences of the *LAT1* gene. This construct was used to replace the *LAT1* locus of the W1536 5B Δ*gcv3* strain by transformation and selection on hygromycin. The N-terminal lipoyl domain and E1 binding domain of Lat1 constructed from the N-terminal 1-184 amino acids was ligated into the yeast integrative vector YIp349, upstream of the CH-tag (N-Lat1 construct). The construct also retained 400 bp of the proximal *LAT1* upstream region. The plasmid was linearized with Bsu361 and integrated into the *TRP1* locus of the Δ*gcv3*Δ*lat1* strain using a high efficiency transformation method.

### Yeast transformations

Plasmids were transformed into the yeast strains using a one-step transformation method published in Chen et al. 1992 [50]. High efficiency transformations of linearized DNA for genomic integration were carried out as described by Gietz and Woods 2002 [51].

### Yeast growth/spotting assays

For spotting assays, strains were grown for 16h in SCD dropout media with the appropriate selection (-URA or ‒URA ‒TRP), harvested and adjusted to an OD_600_ of 0.5. A dilution series of undiluted, 1:10, 1:100 and 1:1000 was made and 2 μl of each dilution were spotted on SCD-URA, SCG-URA, SCG-URA+100 μM LA, SCG-URA+100 μM C8, YPG, YPG+100 μM LA, and YPG+100 μM C8. The plates were grown at 30°C for 4 days.

### Western blot analysis

Strains were grown in YPG, YPG+ 100μM LA, or 100μM C8 for 24h, after which cells were harvested and total protein samples were prepared by trichloroacetic acid precipitation as described in Platta et al. 2004 [52]. Equivalent amounts of total proteins were separated by SDS-PAGE (12% gel) in order to quantify the steady-state levels of mitochondrial lipoylated enzyme complexes Proteins were transferred to nitrocellulose membrane and decorated with the indicated antisera. Polyclonal rabbit anti-lipoic acid antiserum (EMD Millipore Corporation, USA; RRID AB_212120), was used for detection of lipoylated proteins. Ponceau staining and α-actin (Abcam, UK; AB_449644) immunodetection were used as loading controls. Secondary antibodies were Anti-Mouse IgG (H+L), HRP Conjugate (Promega, USA; RRID AB_430834) or Immun-Star Goat Anti-Rabbit (GAR)-HRP conjugate antibody (Bio-Rad Laboratories, lnc. Hercules USA, RRID: AB_11125757). Slight variations in relative signal strength of Lat1-LA versus Kgd2-LA are likely the result of slight variations in affinity of the different anti-LA antibody batches used.

## DECLARATIONS

### Ethics approval and consent to participate

No animal research; not applicable.

No human subjects; not applicable.

### Consent for publication

No individual person’s data used in study; not applicable.

### Availability of data and materials

All data generated or analyzed during this study are included in this published article and the additional data file.

## Author contributions

LPP planned the experiments, carried out the experimental work, wrote the article and generated the Figures. MTR did additional experimental work and participated in writing of the article. JKH planned experiments and participated in the writing of the article. CLD planned experiments, supervised the work and participated in the writing of the article. AJK planned experiments, supervised the work and participated in the writing of the article.

## Acknowledgements

We thank Telsa Mittelmeier for critical reading of our manuscript and her suggestions. We are grateful to Dr. Alexander Tzagoloff at Columbia University, New York, NY USA, for hosting LPP in his lab for three months and sharing expertise and materials.

