## Additional Data Figures A1 to A15 for "Genetic dissection of the mitochondrial lipoylation pathway in yeast"

Figure A1

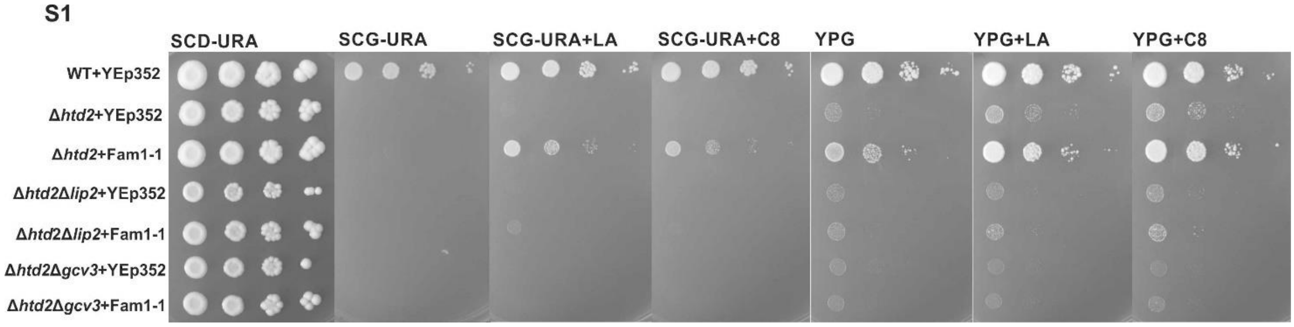


Figure A1. Analysis of growth of lipoylation deficient strains expressing Fam1-1. Extended spotting assay data corresponding to Figure 2A, including growth data of the identical strains on YPG media. Growth assay analysis of wild type+YEp352, Δ*htd2*+YEp352, Δ*htd2*+YEp352mtFam1-1, Δ*htd2*Δ*lip2*+YEp352, Δ*htd2*Δ*lip2*+YEp352mtFam1-1, Δ*htd2*Δ*gcv*3+YEp352 and Δ*htd2*Δ*gcv3*+YEp352mtFam1-1 were spotted on SCD-URA/YPG plates as a general growth control, as well as on SCG-URA/YPG supplemented with LA or C8. Plates were incubated at 30$^{o}$C for 4 days.

Figure A2

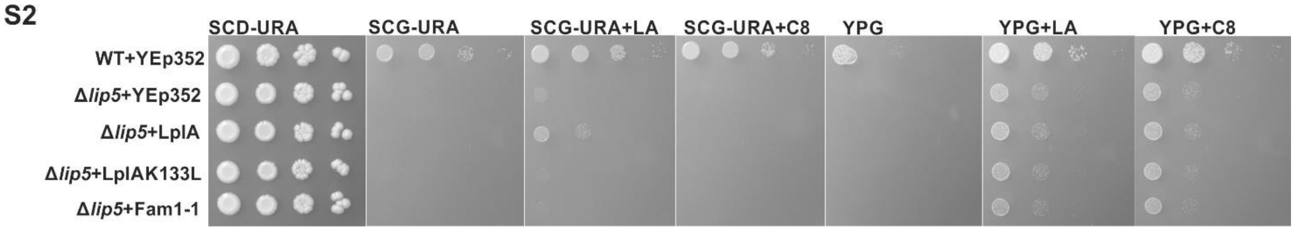


Figure A2. Analysis of growth of the LA synthesis deficient Δ*lip5* strain expressing LplA. Extended spotting assay data corresponding to Figure 4A, including growth data of the identical strains on YPG media. Growth assay of Δ*lip5* strain complemented with YEp352-LplA and YEp352-LplAK133L (negative control) on SCD-URA/YPG plates as a general growth control, as well as on SCG-URA/YPG supplemented with LA or C8. Plates were incubated at 30$^{o}$C for 4 days.

Figure A3

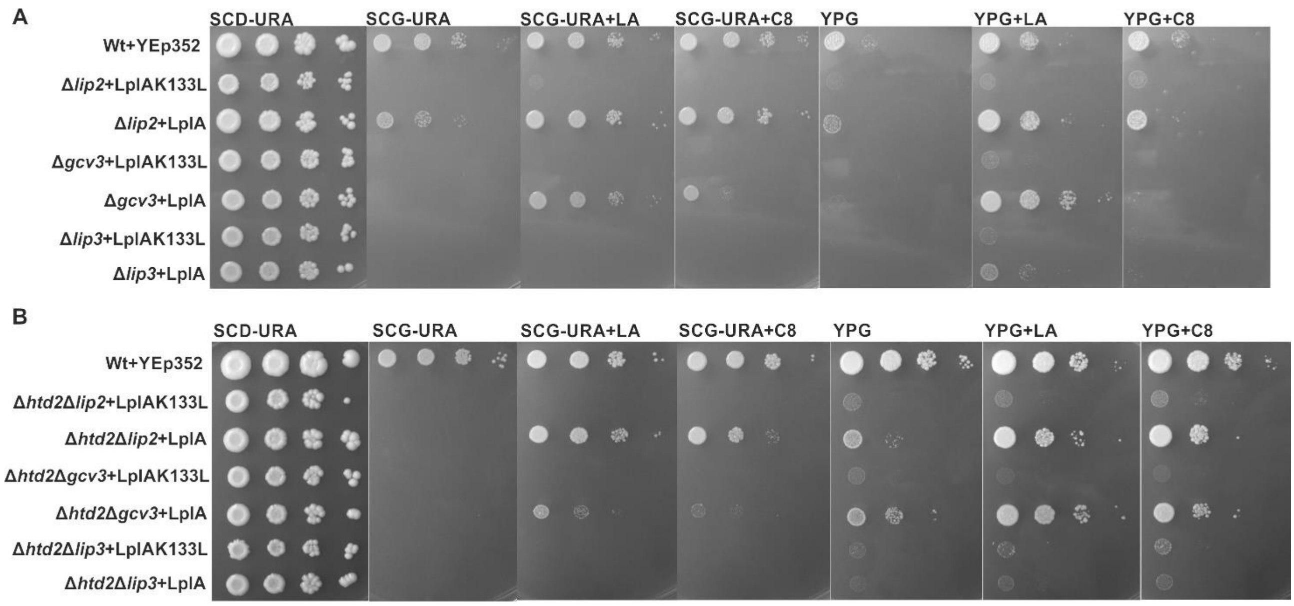


Figure A3. Growth assay analysis of lipoylation deficient strains complemented with LplA. Extended spotting assay data corresponding to Figure 5, including growth data of the identical strains on YPG media. A) Δ*lip2*, Δ*gcv3* and Δ*lip3* complemented with YEp352-LplA as well as YEp352-LplAK133L as a negative control were spotted on SCD-URA/YPG plates as a general growth control, as well as on SCG-URA/YPG supplemented with LA or C8. Plates were incubated at 30$^{o}$C for 4 days.

(B) Δ*htd2*Δ*lip2*, Δ*htd2*Δ*gcv3* and Δ*htd2*Δ*lip3* complemented with YEp352-LplA as well as YEp352-LplAK133L as a negative control were spotted on SCD-URA/YPG plates as a general growth control, as well as on SCG-URA/YPG supplemented with LA or C8. Plates were incubated at 30$^{o}$C for 4 days.

Figure A4

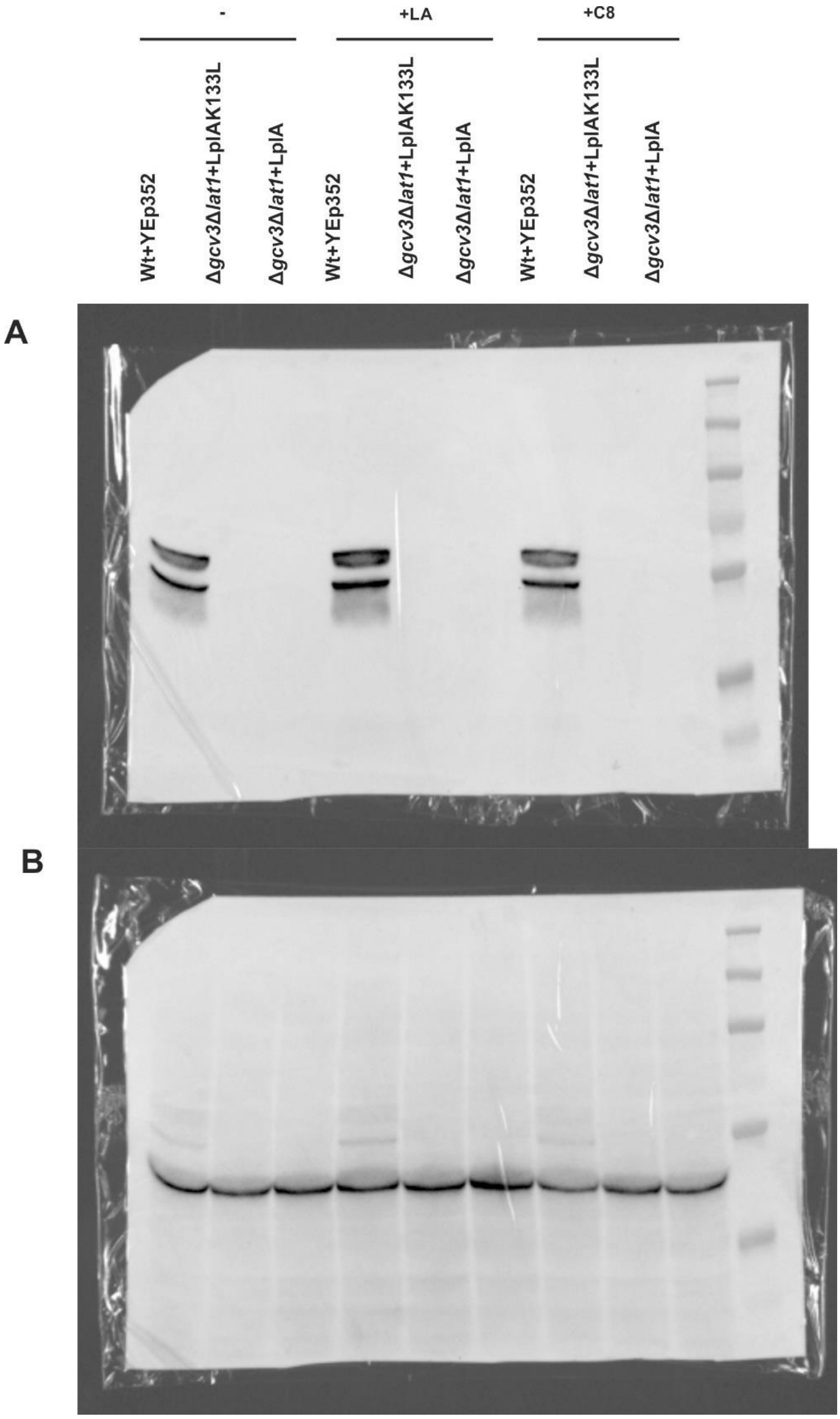


Figure A4. Original image files used to compose Figure 7 B. A) Original anti-LA signal. B) Original anti-actin signal. A shadow of the LA-signal matching the image on panel A is visible.

Figure A5

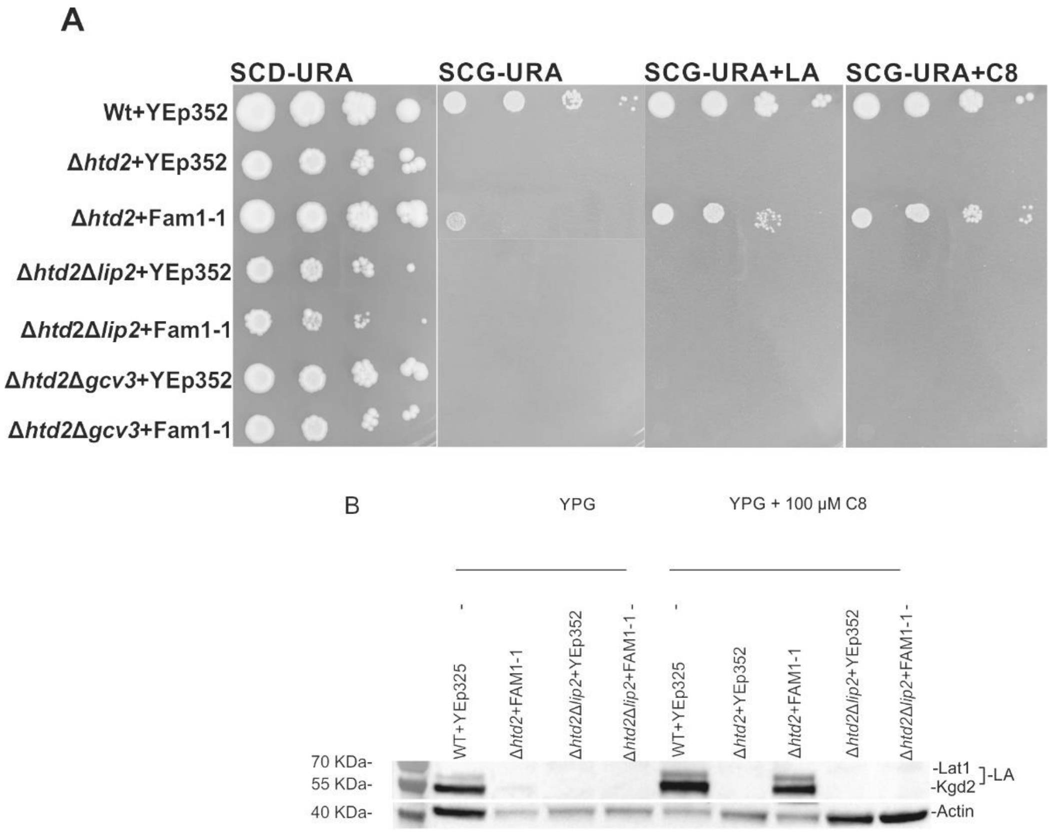


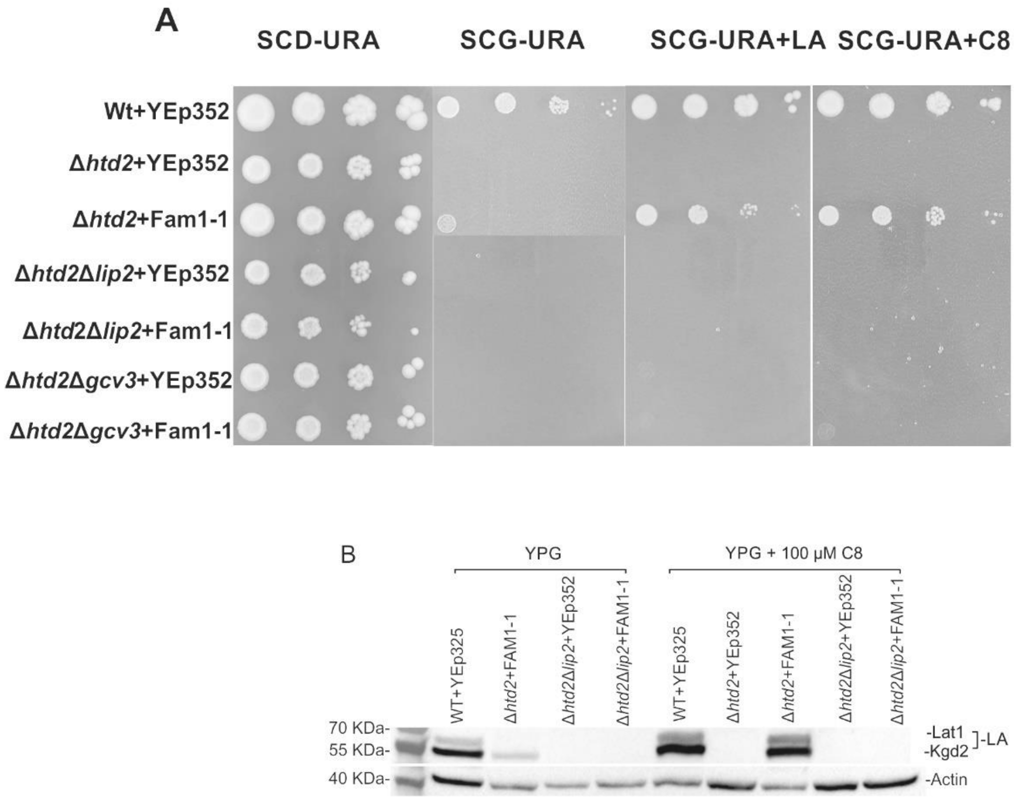


Figure A5 repeat experiments for Figure 2 in main manuscript

Figure A6

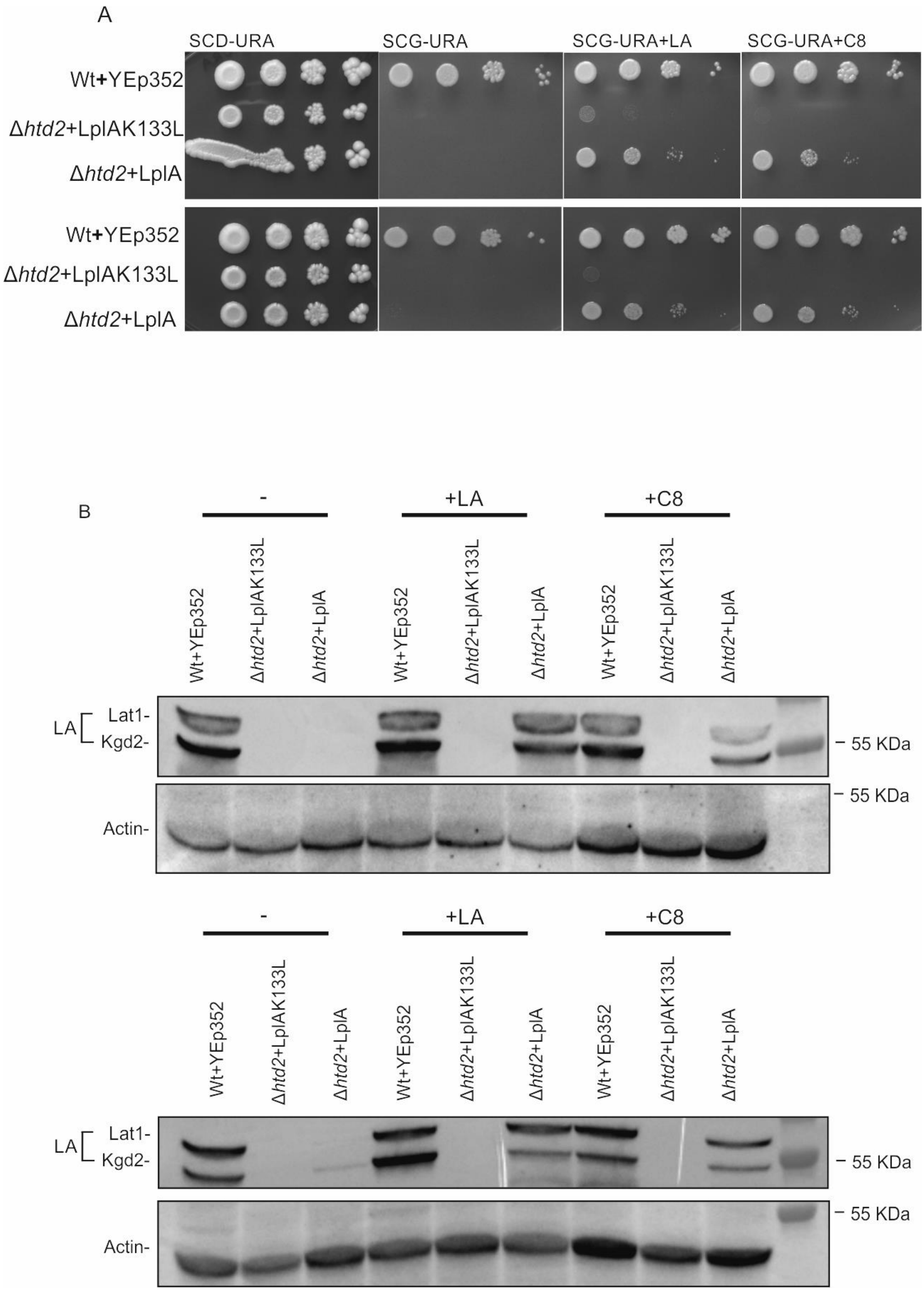


Figure A6. Repeats of experiments depicted in Figure 3 of the main document

Figure A7

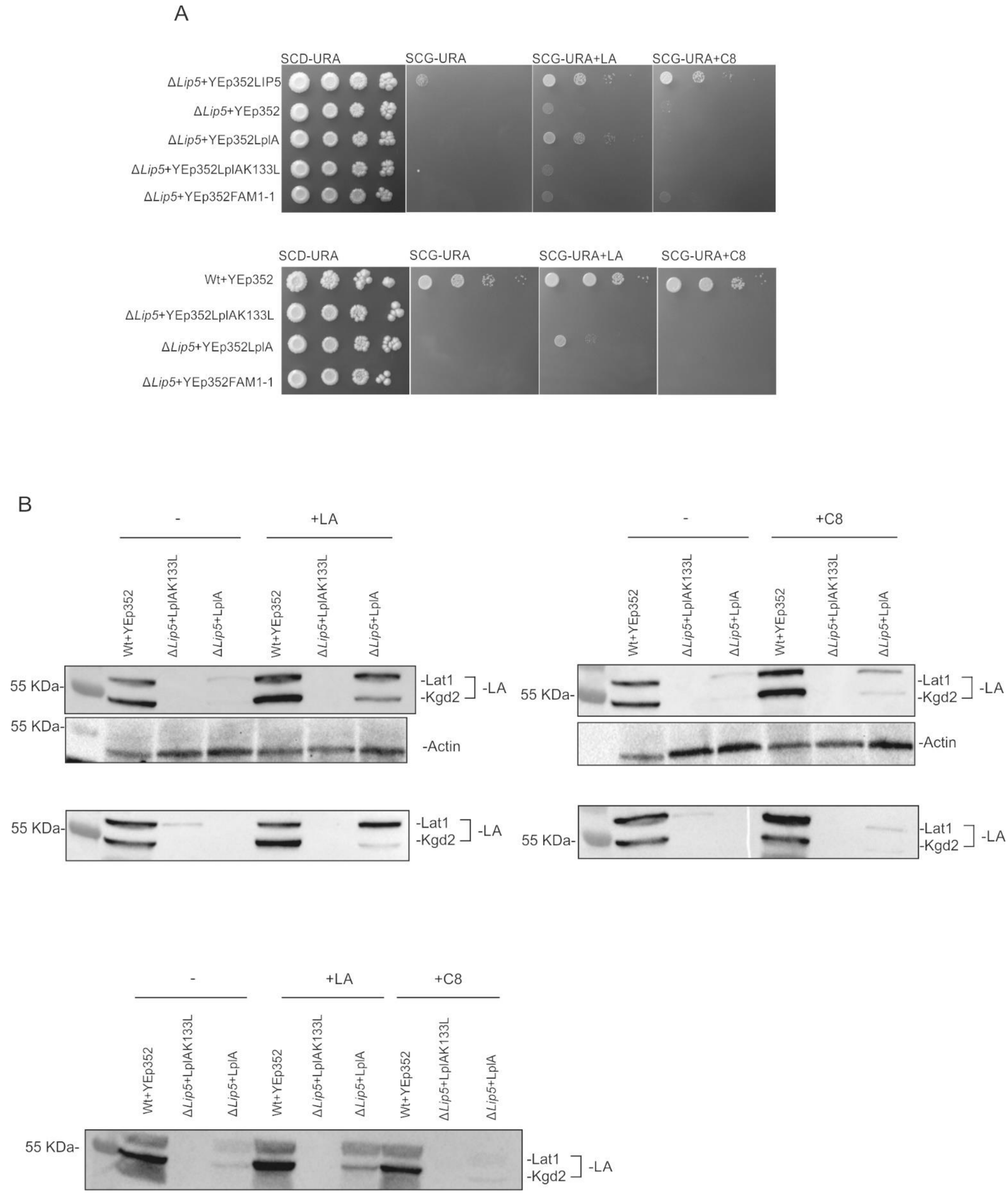


Figure A7. Repeats of experiments depicted in Figure 4 of the main document

Figure A8

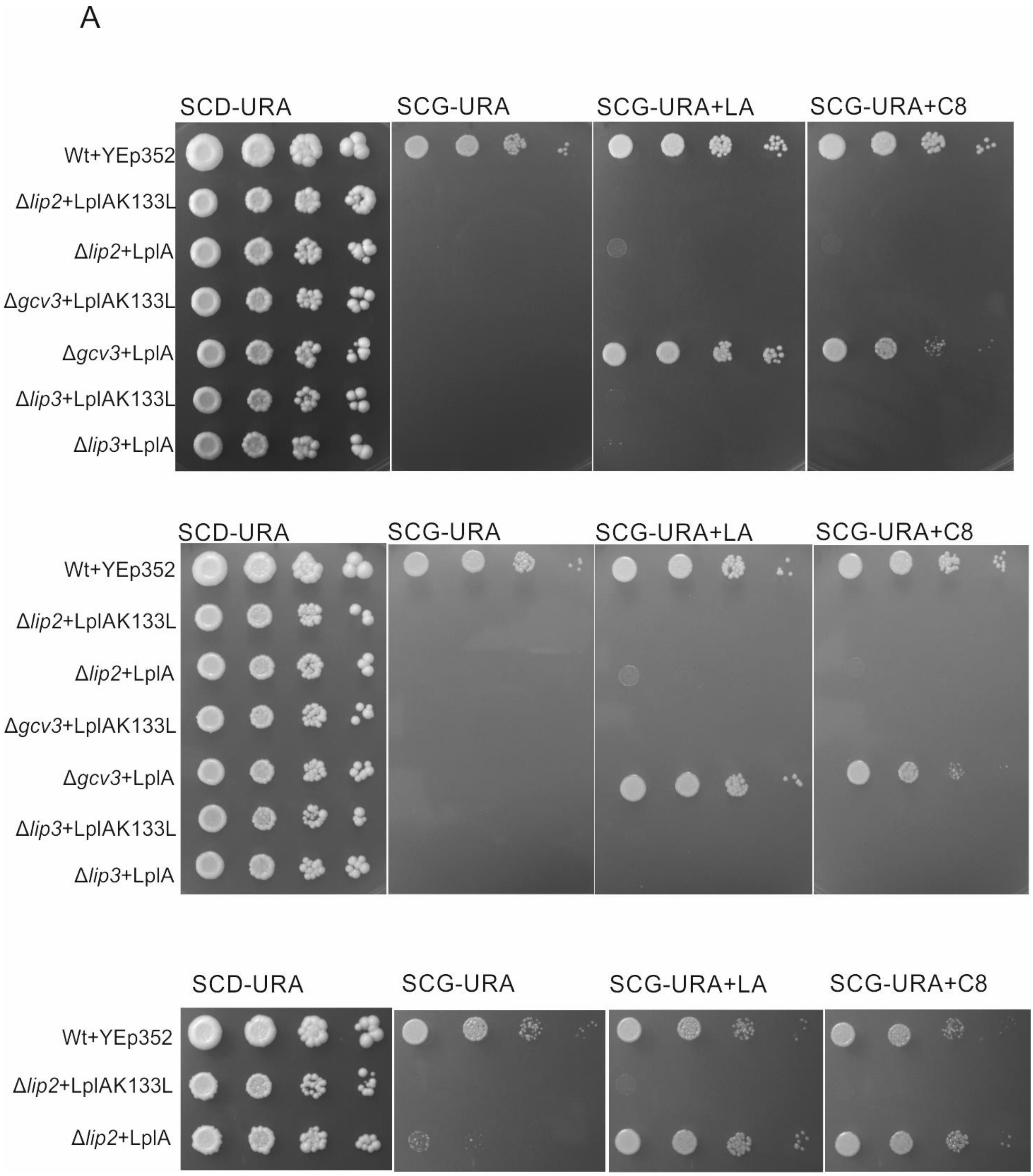


Figure A8. Repeats of Experiments depicted in Figure 5 of the main document

Figure A9

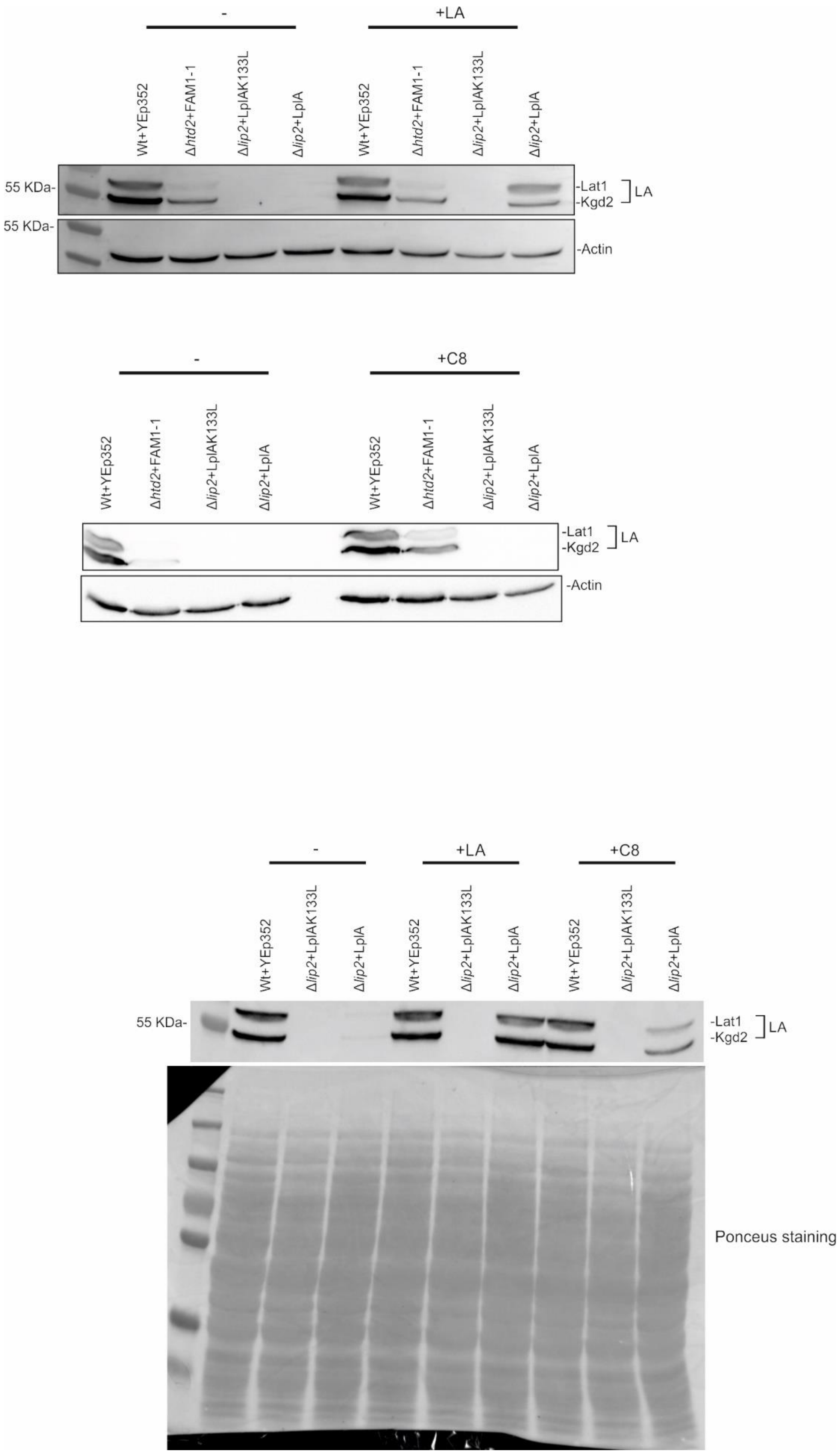


Figure A9. Repeats of Experiments depicted in Figure 6A of the main document

Figure A10

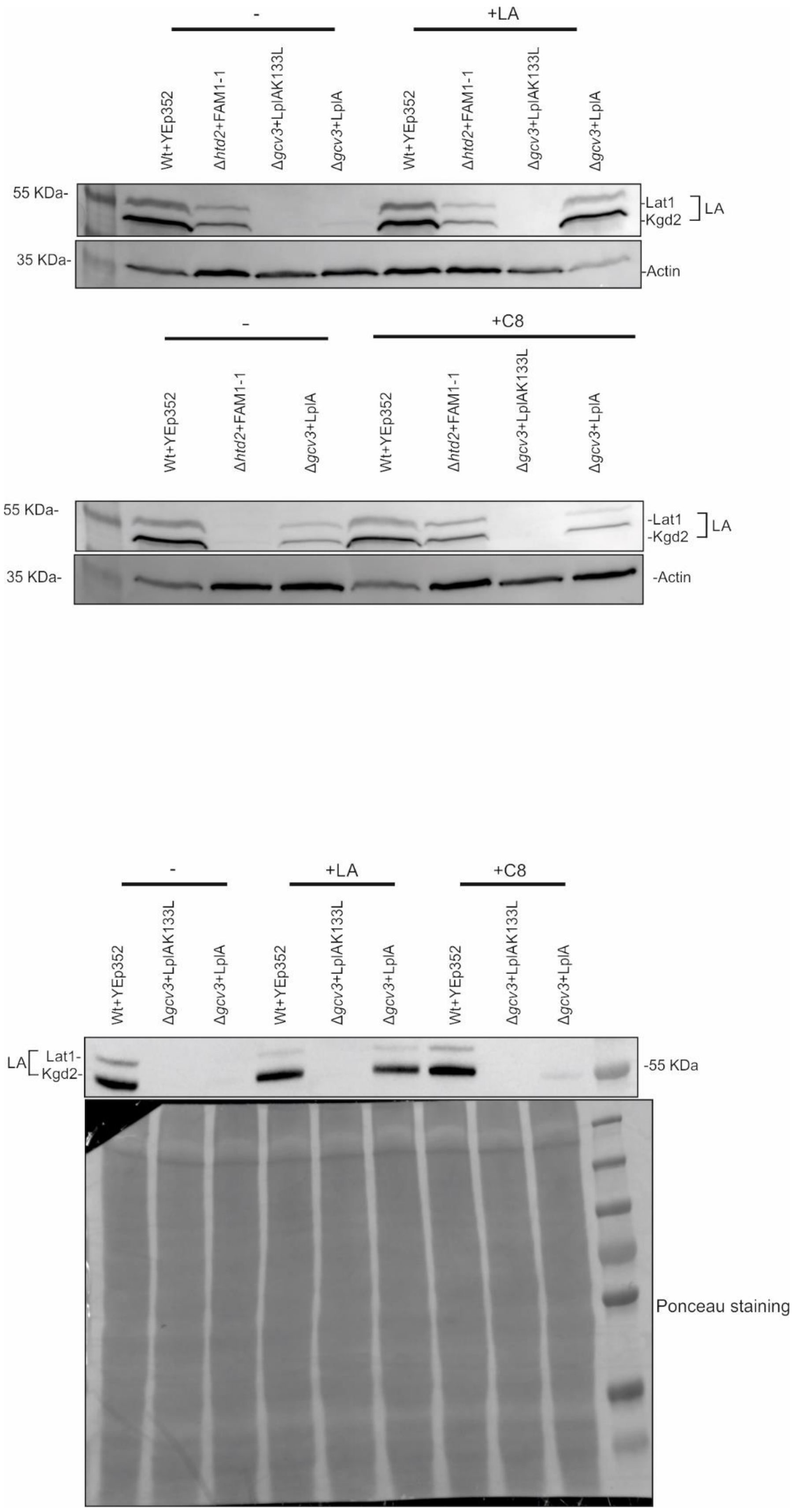


Figure A10. Repeats of Experiments depicted in Figure 6B of the document

Figure A11

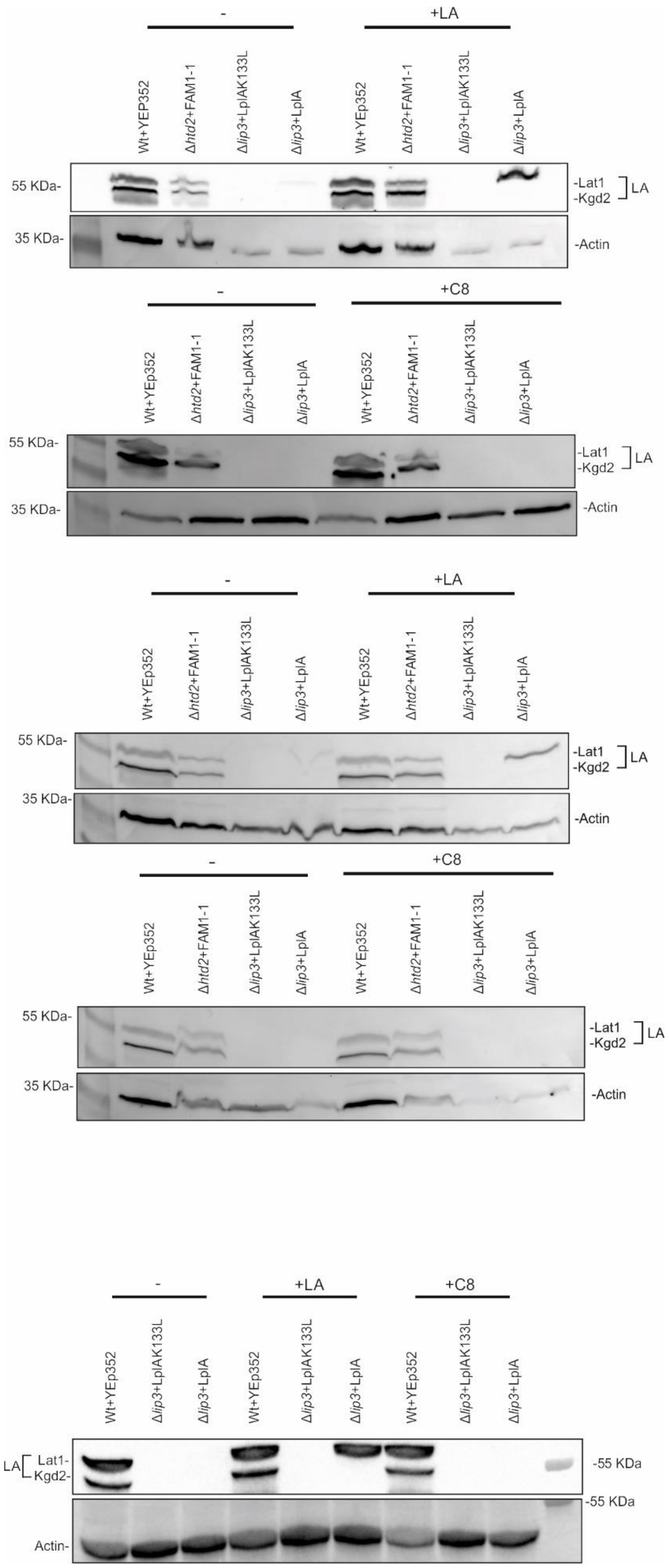


Figure A11. Repeats of Experiments depicted in Figure 6C of the document

Figure A12

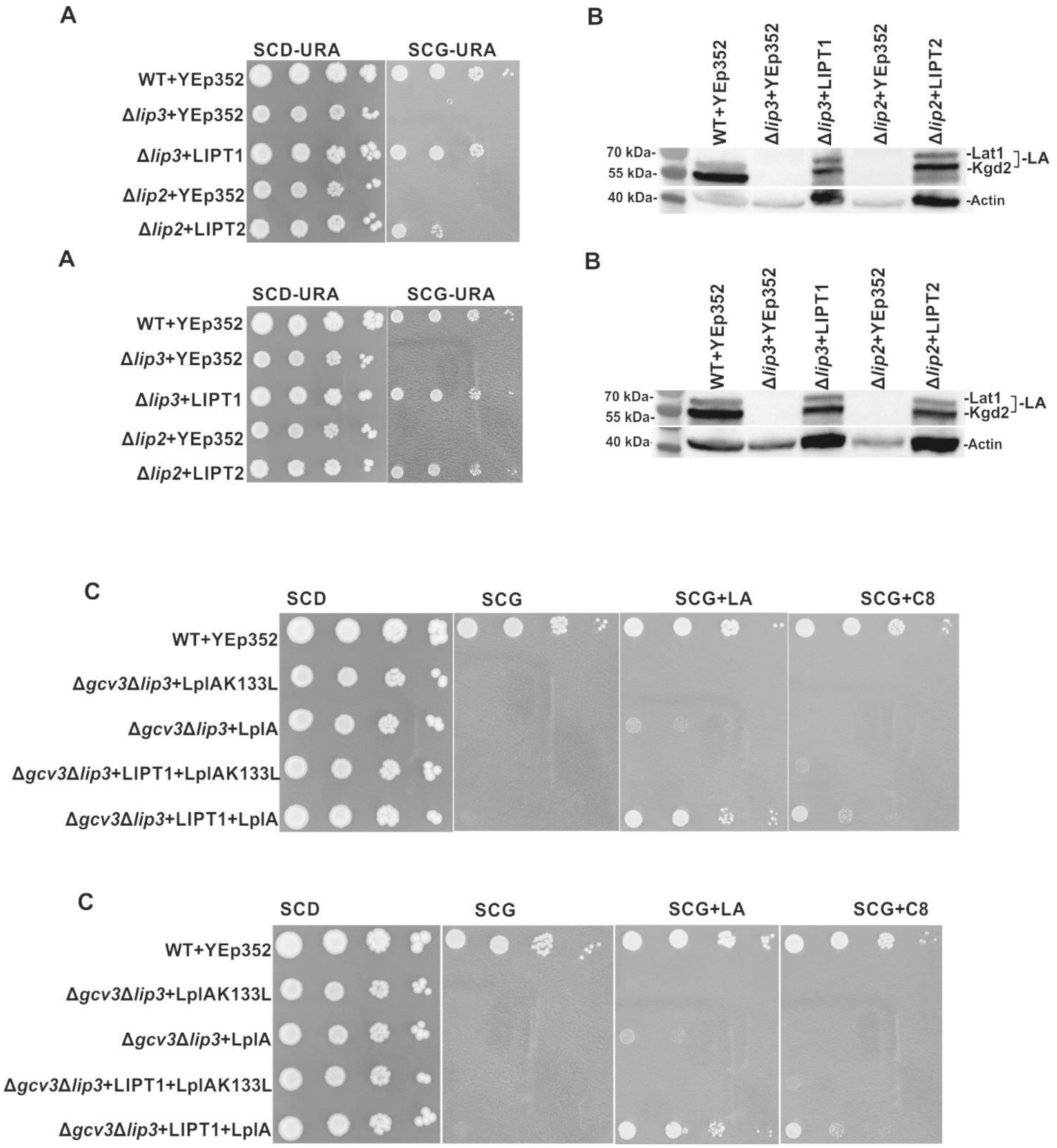


Figure A12. Repeats of Experiments depicted in Figure 8 of the document

Figure A13 A. C8 titration growth assay (three independent repeats)

Repeat#1

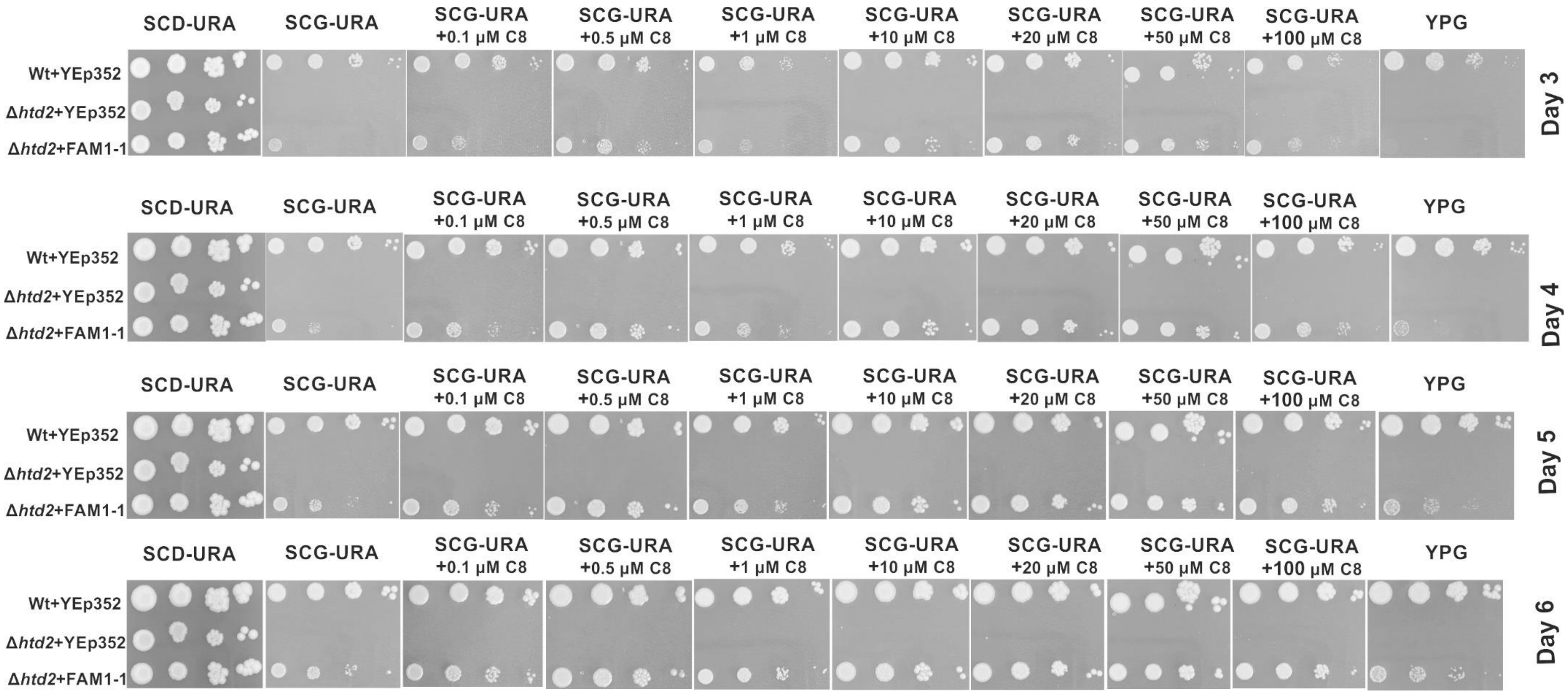


Figure A13 A: Growth assays of wild type +YEp352, Δ*htd2*+YEp352 and Δ*htd2*+YEp352mtFam1-1. The strains were grown to logarithmic growth phase, harvested and adjusted to $OD_{600}$ of 0.5. A dilution series of undiluted, 1:10, 1:100, 1:1000 was made and 2µl of cells of each dilutions were spotted on a SCD-URA plate as a general growth control, on SCG-URA media supplemented with C8 (0.1µM, 0.5µM, 1µM, 10µM, 20µM, 50µM and 100µM) and on YPG. Plates were incubated at 30$^{0}$C for 6 days and the growth was observed from day 3 to day 6. At 1 µM and 100 µM C8 concentrations, we see a slight inhibition in growth. We attribute this to increased ethanol concentration in the growth media (C8 stock concentrations 10 mM and 100mM in 70 % ethanol, respectively).

Figure A13 B

Repeat#2

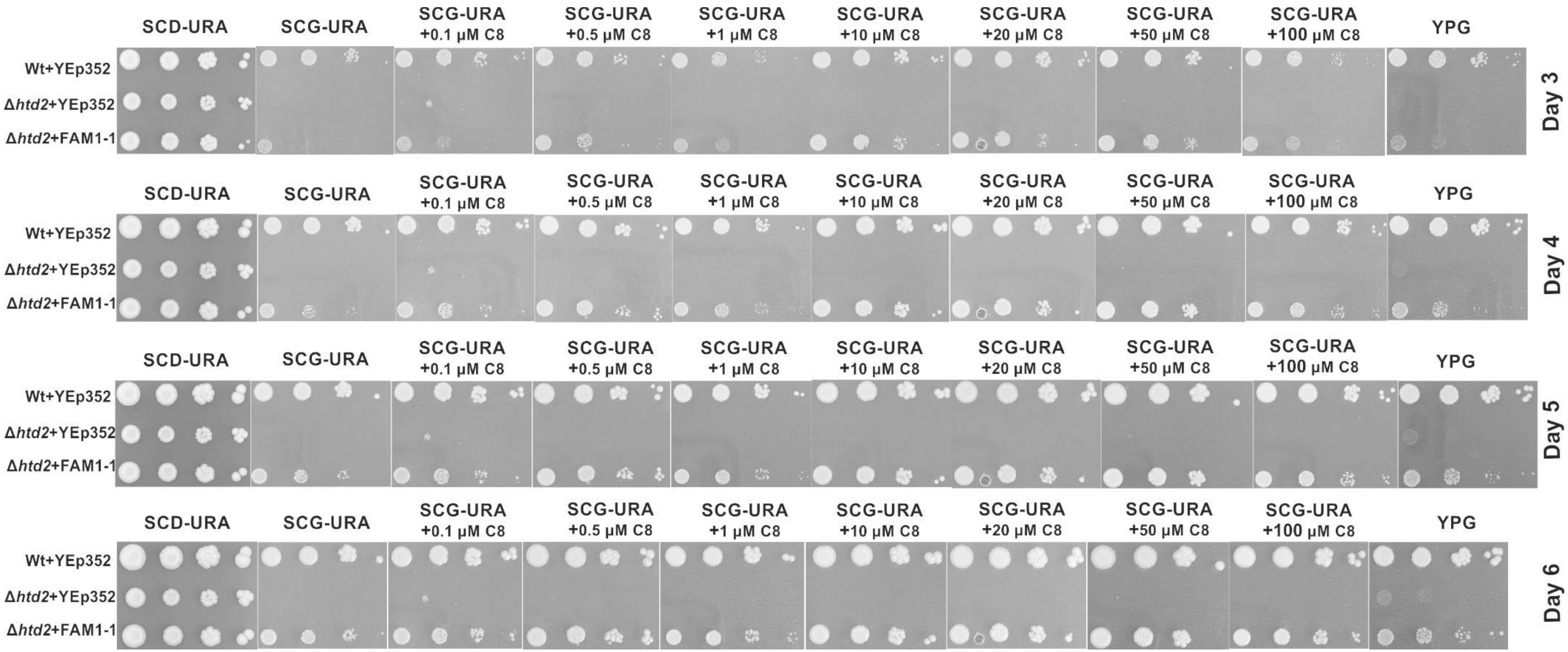


Figure A13 B: Growth assays of wild type +YEp352, Δ*htd2*+YEp352 and Δ*htd2*+YEp352mtFam1-1. The strains were grown to logarithmic growth phase, harvested and adjusted to $OD_{600}$ of 0.5. A dilution series of undiluted, 1:10, 1:100, 1:1000 was made and 2µl of cells of each dilutions were spotted on a SCD-URA plate as a general growth control, on SCG-URA media supplemented with C8 (0.1µM, 0.5µM, 1µM, 10µM, 20µM, 50µM and 100µM) and on YPG. Plates were incubated at 30$^{o}$C for 6 days and the growth was observed from day 3 to day 6. At 1 µM and 100 µM C8 concentrations, we see a slight inhibition in growth. We attribute this to increased ethanol concentration in the growth media (C8 stock concentrations 10 mM and 100mM in 70 % ethanol, respectively).

Figure A13 C

Repeat #3

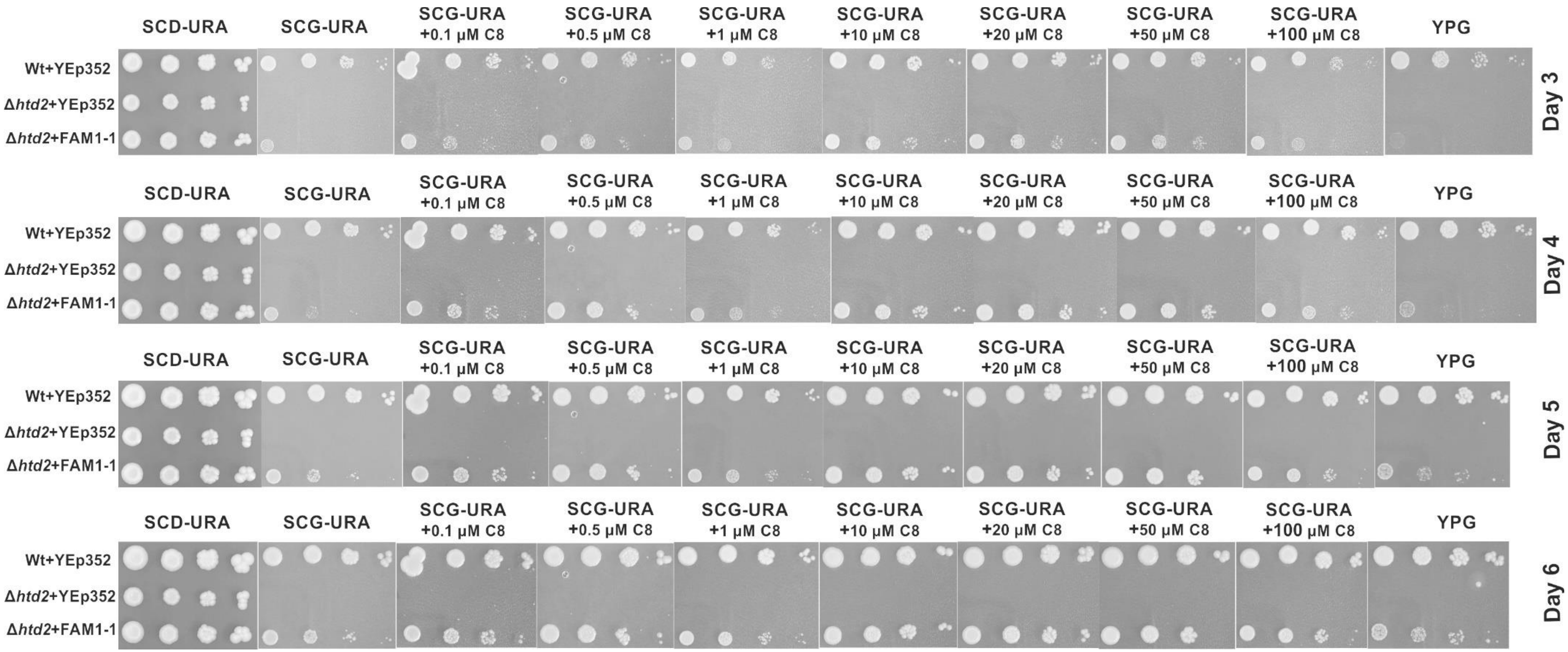


Figure A13 C: Growth assays of wild type +YEp352, Δ*htd2*+YEp352 and Δ*htd2*+YEp352mtFam1-1. The strains were grown to logarithmic growth phase, harvested and adjusted to $OD_{600}$ of 0.5. A dilution series of undiluted, 1:10, 1:100, 1:1000 was made and 2µl of cells of each dilutions were spotted on a SCD-URA plate as a general growth control, on SCG-URA media supplemented with C8 (0.1µM, 0.5µM, 1µM, 10µM, 20µM, 50µM and 100µM) and on YPG. Plates were incubated at $30^{o}$C for 6 days and the growth was observed from day 3 to day 6. At 1 µM and 100 µM C8 concentrations, we see a slight inhibition in growth. We attribute this to increased ethanol concentration in the growth media (C8 stock concentrations 10 mM and 100mM in 70 % ethanol, respectively).

Figure A14. C8 titration lipoic acid western blotting

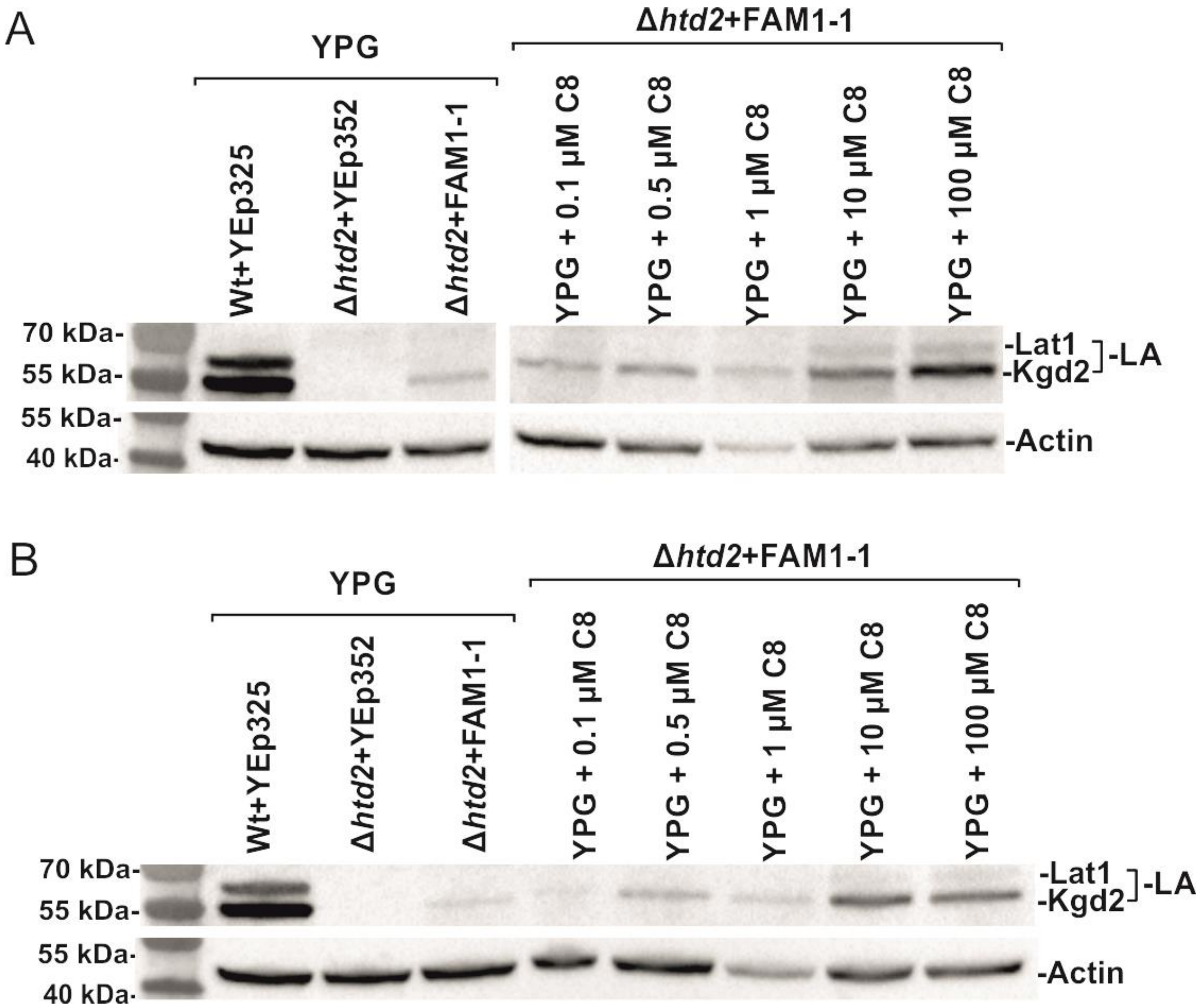


Figure A14: Western blot analysis of extracts of mtFam1-1 complemented strains. Whole cell extracts were collected from cells expressing YEp352mtFam1-1 or YEp352 as a negative control, after 24 h of growth on YPG media at 30°C. Whole cell extracts from wild type+YEp352, Δhtd2+YEp352 and Δhtd2+YEp352mtFam1-1 grown without supplements (YPG) or with 0.1µM, 0.5µM, 1µM, 10µM and 100µM C8 supplementation. Analysis was done by probing with anti-LA serum and anti-actin serum as a loading control. PageRulerTM Prestained protein ladder was used as marker.

Figure A15 A. C8 titration lipoic acid western blotting (A-C three independent repeats)

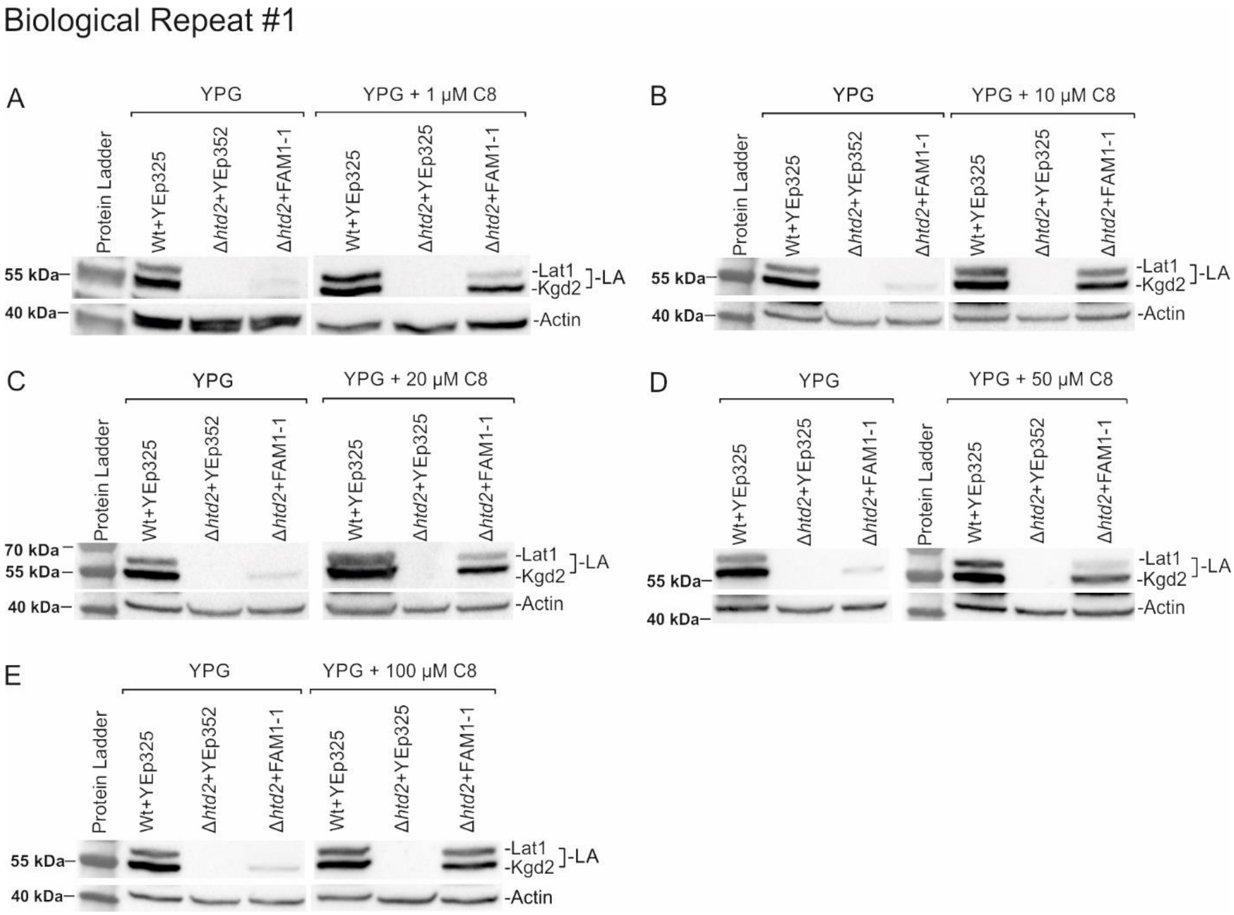


Figure A15 A: Western blot analysis of extracts of mtFam1-1 complemented strains. Whole cell extracts were collected from cells expressing YEp352mtFam1-1 or YEp352 as a negative control, after 24 h of growth on YPG media at 30$^{o}$C. Whole cell extracts from wild type+YEp352, Δhtd2+YEp352 and Δhtd2+YEp352mtFam1-1 grown without supplements (YPG) or with 1µM (A), 10 µM (B), 20 µM (C), 50 µM (D) and 100µM (E) C8 supplementation. Analysis was done by probing with anti-LA serum and anti-actin serum as a loading control. PageRulerTM Prestained protein ladder was used as marker.

Figure A15 B

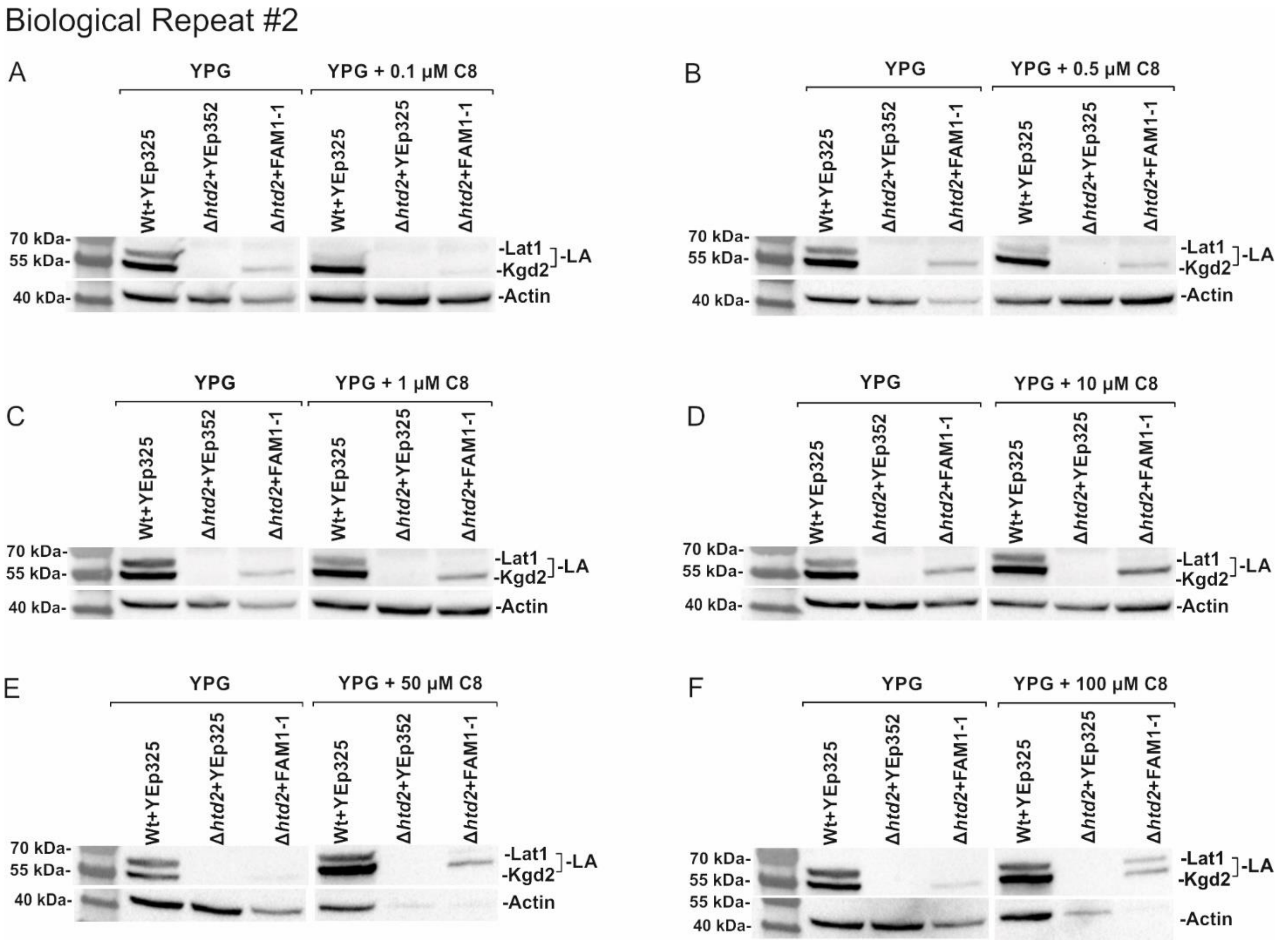


Figure A15 B: Western blot analysis of extracts of mtFam1-1 complemented strains. Whole cell extracts were collected from cells expressing YEp352mtFam1-1 or YEp352 as a negative control, after 24 h of growth on YPG media at 30$^{o}$C. Whole cell extracts from wild type+YEp352, Δhtd2+YEp352 and Δhtd2+YEp352mtFam1-1 grown without supplements (YPG) or with 0.1µM (A), 0.5 µM (B), 1 µM (C), 10 µM (D), 50µM (E) and 100 µM (F) C8 supplementation. Analysis was done by probing with anti-LA serum and anti-actin serum as a loading control. PageRulerTM Prestained protein ladder was used as marker.

Figure A15 C

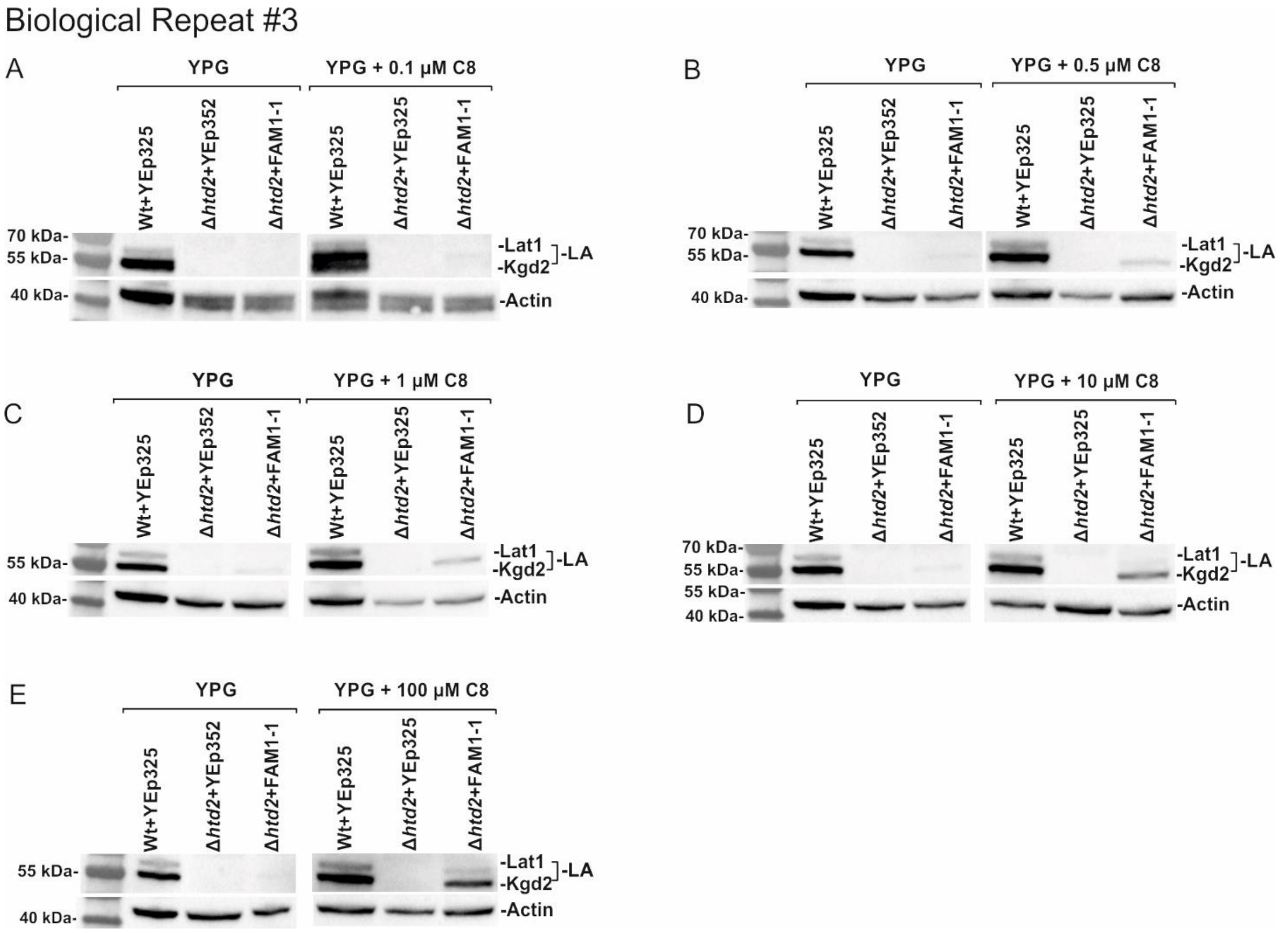


Figure A15 C: Western blot analysis of extracts of mtFam1-1 complemented strains. Whole cell extracts were collected from cells expressing YEp352mtFam1-1 or YEp352 as a negative control, after 24 h of growth on YPG media at 30$^{o}$C. Whole cell extracts from wild type+YEp352, Δhtd2+YEp352 and Δhtd2+YEp352mtFam1-1 grown without supplements (YPG) or with 0.1µM (A), 0.5 µM (B), 1 µM (C), 10 µM (D) and 100 µM (E) C8 supplementation. Analysis was done by probing with anti-LA serum and anti-actin serum as a loading control. PageRulerTM Prestained protein ladder was used as marker.
